# Genomic signatures of adaptation to abiotic stress from a geographically diverse collection of chile peppers (Capsicum spp.) from Mexico

**DOI:** 10.1101/2023.08.13.553093

**Authors:** Vivian Bernau, Michael Kantar, Lev Jardon Barbolla, Jack McCoy, Kristin L. Mercer, Leah K. McHale

## Abstract

Extreme environmental variability requires the identification of genetic diversity that can help crops withstand difficult abiotic conditions. Understanding the genetic basis of adaptation to abiotic stress can provide tools for adapting agriculture to changing climates. Crop landraces and their wild ancestors from centers of domestication have often evolved across diverse environmental conditions and provide a unique opportunity to locate loci harboring alleles that could contribute to abiotic stress tolerance, among other traits. We collected chile peppers (*Capsicum* spp.) from farmers in southern Mexico along environmental transects crossing temperature, precipitation, and elevational gradients. Using 32,623 filtered SNPs generated from genotyping-by-sequencing (GBS), we conducted an environmental association analysis (EAA) combined with two outlier analyses, F_ST_ and spatial ancestry analyses to detect priority candidate loci associated with climate and soil phenotypes relevant to drought tolerance. Even though cultivated species may be shielded from some environmental selection by management practices (e.g., irrigation), we found fifteen priority loci whose genetic variation covaried with environmental variation in our EAA and were also allele frequency outliers (i.e., Fst and/or SPA). Thirty-three percent of the priority loci were validated with phenotype in response to water deficit. The haplotype blocks associated with these loci identify unique genetic variants worthy of conservation and harbor genes with abiotic stress-related functions. This work provides valuable information to explore quantitative trait loci (QTLs) underlying abiotic stress tolerance in chile pepper and is a new resource for improving plant breeding around the world.

**Article Summary:** The extreme environmental variability faced by crops requires the identification of potentially adaptive loci for abiotic stressors. We conducted an environmental association analysis (EAA) environments using chile peppers collected in southern Mexico along environmental transects crossing temperature, precipitation, and elevational gradients. We combined EAA with outlier analyses F_ST_ and spatial ancestry analysis to detect priority candidate loci associated with climate and soil phenotypes relevant to drought tolerance that may putatively contribute to abiotic stress adaptation.

## Introduction

Intensifying environmental change has escalated the global push to understand the complex genetic basis of abiotic stress tolerance in plants. Landscape genomics studies are being used to identify patterns of adaptive genetic variation (Manel & Holderegger, 2013; Rellstab et al., 2015; Waldvogel et al., 2020). Landscape genomic methods work on the assumption that environmental variables have provided selective pressure on the genome and can include genome-wide calculations of population statistics and identification of outlier loci (Lotterhos & Whitlock, 2015; Matthey□Doret & Whitlock, 2019) as well as environmental association analyses (EAA, also known as genome-environment associations—GEA), which identifies adaptive loci by associating genome-wide markers with variables describing the source environment.

EAAs have been applied and tested in model species—*Arabidopsis*, tree species, vertebrates, fungi and several crops of global importance. Adaptive loci identified in *Arabidopsis* were shown to effectively predict relative fitness in a common garden experiment across wide-ranging environments (Fournier-Level et al., 2011; Hancock et al., 2011; also see Alonso-Blanco et al., 2016). EAA studies on sorghum (*Sorghum bicolor* (L.) Moench), maize (*Zea mays* subsp. *mays* L.), barley (*Hordeum vulgare* L.) and foxtail millet (*Setaria italica*) have provided insights into the genetic architecture and genes underlying adaptation (Lasky et al., 2015; C. Li et al., 2021; Romero Navarro et al., 2017; Russell et al., 2016; J. Wang et al., 2020). Furthermore, EAAs of locally adapted soybean landraces (*Glycine max*) and soybean’s wild progenitor, *Glycine soja* L., have identified dozens of interesting genomic regions associated with environmental variables (Y. Li et al., 2020). So far, EAA has not been conducted with germplasm from the genus *Capsicum*, or any other horticultural crop. Furthermore, examples of EAA in crop plants which are paired with experimental phenotypic data to test if the same loci identified by environmental phenotypes are also identified by measured phenotypes are more limited (Fournier-Level et al., 2011; Hancock et al., 2011; Lasky et al., 2015; C. Li et al., 2021; Y. Li et al., 2020).

There are a variety of statistical models and software available for this type of “reverse ecology” (reviewed in Rellstab et al., 2015). Genome wide association studies (GWAS) provide a computationally efficient method which is commonly used to identify genes controlling variation in target traits by making associations between single nucleotide polymorphisms and measured phenotypes. GWAS methods have been employed in crop plants to identify the genetic architecture of complex traits (Saini et al., 2021; Sukumaran et al., 2015; Zhao et al., 2011), including those related to drought tolerance (Adewale et al., 2018; Morris et al., 2013; N. Wang et al., 2016; Xue et al., 2013). However, one drawback of GWAS is that identified loci only explain a subset of the total genetically controlled variation. Similarly, GWAS models, especially those employing a relatively low number of individuals, have traditionally been ineffective at identifying rare alleles or common, small effect alleles because of the complex genetic structure that often underlies crop plants. However, more recent GWAS models with improved statistical power are beginning to break this barrier (Liu et al., 2016; Miao et al., 2018).

Furthermore, GWAS can be combined with other landscape genomic approaches to identify regions of the genome that have undergone selection and may therefore include climate adaptive loci. Methods identifying signatures of selection are particularly useful in exploring natural populations (Pardo-Diaz et al., 2015; Schlötterer, 2003; Stinchcombe & Hoekstra, 2008) and should be less biased towards large-effect loci than most GWAS approaches (Martin & Jiggins, 2013; Pardo-Diaz et al., 2015). Outlier tests based on F_ST_ (the fixation index) are the most common way to identify locus-specific population differentiation; they detect greater differences in allele frequency than expected under neutral evolution (Günther & Coop, 2013; Weir & Cockerham, 1984). Furthermore, signatures of selection can be investigated within spatial and environmental contexts to identify important loci with abrupt changes in frequency across the landscape or environmental clines (Coop et al., 2010; De Mita et al., 2013; Joost et al., 2008; Yang et al., 2012). By detecting regions of the genome under selection, interpopulation measures such as *F_ST_* and Spatial Ancestry (SPA) analyses can be used to identify loci that may contribute to local adaptation.

Local adaptation occurs when a population evolves to become more suited to its environment due to selection pressures. However, local adaptation can be impeded by high rates of gene flow between populations and large population sizes (Lenormand, 2002; Slatkin, 1973). Furthermore, human selection in domesticated crops (e.g., by farmers for taste, color, or harvest characteristics) can counter or corroborate environmental selection pressures. Nevertheless, repeated selection within one environment, such as a single farmer’s parcel of land, could contribute to local adaptation and adaptation to incremental climate change (Mercer et al., 2008). Thus, crop wild relatives and landraces—traditional varieties of crops conserved and cultivated by farmers for generations—provide a unique resource to study local adaptation. Genes underlying local adaptation are of great interest in crop improvement, as variation at these loci can be harnessed to help other crop populations and improved crop varieties perform better in harsh environments (Corrado & Rao, 2017).

Abiotic stress tolerance has a complex genetic basis, but many stress responsive genes are related to abscisic acid (ABA) biosynthesis and signaling. In pepper, the genes *CaDIK1* and *CaDIL1* are involved in enhanced stomatal control as a response to ABA, the production of which is triggered by drought stress (Lim et al., 2018, 2020). However, the expression of these genes is only a small part of the ABA pathway contributing to a drought tolerant phenotype, and further work to understand signaling pathways is essential to understanding adaptive responses to abiotic stress. EAA may be helpful in identifying further candidate genes related to ABA biosynthesis and signaling, thus conferring abiotic stress tolerance.

In this study, we examine environmental and phenotypic associations with genomic data to identify putative abiotic stress tolerance loci of adaptive interest in wild and landrace chiles collected across varied landscapes in a crop center of diversity in southern Mexico. Our specific objectives were to identify DNA sequence variation 1) associated with high resolution environmental “phenotypes” obtained from global datasets using EAA, 2) under selection between populations (through *F_ST_*), and 3) under selection across the landscape (using SPA). Finally, in order to provide a further comparison and support for loci found in objectives 1-3, we 4) associated DNA sequence variation with measured phenotypes from a greenhouse experiment manipulating drought. Our list of candidate loci could provide fodder for further elucidation of mechanisms underlying abiotic stress tolerance in this era of climate change.

## Materials and Methods

DNA from 190 lines generated from 103 accessions of pepper (*Capsicum* sp.) collected in the Mexican states of Oaxaca and Yucatan across environmental and elevational gradients was used to create DNA libraries and pools of 48 genotypes were sequenced using genotype by sequencing (GBS with NextSeq and HiSeq 2500 platforms). The resulting sequence data was processed per “Mexican Collection” in Taitano et al. (2018) with the TASSEL GBS Pipeline 5.2.3 (Glaubitz et al., 2014) and the *C. annuum* cv. CM334 reference genome to align with Bowtie2 (Langmead & Salzberg, 2012), which identified a total of 32,623 filtered SNPs.

By querying two global datasets [WorldClim2 (Fick & Hijmans, 2017) and ISRIC (World Soil Information database; Hengl et al., 2017)] using latitude and longitude from the original collections, we amassed nineteen bioclimatic variables describing variation in temperature and precipitation at various time scales and seven edaphic variables describing topsoil (0-30 cm) and subsoil (30-200cm). We calculated other metrics, such as monthly potential evapotranspiration (PET) (Thornthwaite and Mather, 1955; Thornthwaite et al., 1957) and drought index (DI) using precipitation (P) data:

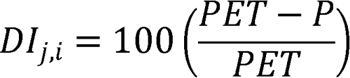

All variables were scaled to a mean of zero and a standard deviation of one, and a principal components analysis (PCA) was conducted. Pearson correlation coefficients were also calculated (Figures A & B in File S1). A summary of bioclimatic and edaphic variables tested can be found in Supplemental Table 1 in File S2.

An F_ST_ outlier scan with six genetic clusters [from fastSTRUCTURE analysis; (Raj et al., 2014; Taitano et al., 2019)] identified the top 100 loci (above 99.7 percentile) with differential allele frequency across groups based on theta (θ), the variance-based *F_ST_*estimate (Weir and Cockerham (1984). Spatial ancestry analysis (SPA) was conducted to detect loci showing steep gradients in allele frequency across space (>99.9^th^ percentile) that were likely the products of a combination of selection and isolation by distance (Yang et al., 2012).

EAA identified associations between SNPs and environmental variables using three methods: a general linear model (GLM), a mixed linear model (MLM), and FarmCPU, a multi-locus mixed model (MLMM) (Yin et al., 2017). We controlled false-positives (type I error) by incorporating kinship matrices and principal components to account for population structure (Kang et al., 2008; Yin et al., 2017; Zhang et al., 2010) and covariates of elevation, latitude, and longitude to account for spatial structure. Since the environmental phenotypes were not normally distributed, we estimated a separate p-value significance threshold for each variable (based on the 95% quantile value of the minimum p-value of 1000 permutations), which ranged from 5.45 × 10^-12^ to 3.26 × 10^-5^. To estimate the percent variance explained (PVE), we used the SNP for each association with the lowest p-value in each haplotype block using a comparison of the linear model’s R^2^ with and without the locus in Tassel (v5.0) (Bradbury et al., 2007; Glaubitz et al., 2014). The haplotype blocks were estimated ignoring markers with minor allele frequency (MAF) < 0.05 with 95% confidence bounds on *D*′, the normalized coefficient of linkage disequilibrium; blocks identified have 95% of informative comparisons in “strong LD”.

To provide additional priority loci to the list from *F_ST_*, SPA, and EAA, we performed a phenotypic genome-wide association study (GWAS). Phenotypic trait data was collected from plants grown in a replicated greenhouse experiment where all chile pepper lines experienced either well-watered or water deficit conditions (the latter imposed by restricting water to 1/3 of that supplied to well-watered pots). Traits included area of the upper-most, fully expanded leaf and its dry biomass, specific leaf area (SLA; the ratio of leaf area to dry mass), severity of aborted leaves (1-5 scale), fruiting status (0 or 1), number of fruits, above-ground biomass (partitioned into stems, leaves, and fruit), plant height and number of branching nodes. Only plants that fruited were included in analyses for number and biomass of fruits. We analyzed a general linear mixed model testing the fixed effects of line, water treatment, and their interaction and the random effects of blocks over time and bench nested within block. BLUPs (best linear unbiased predictors) of traits were calculated for the 156 lines under each water treatment and used to calculate three estimates of trait stability under drought:

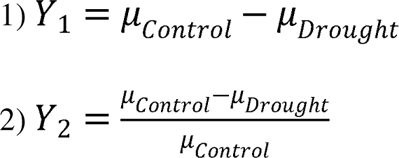

BLUPs were analyzed in the GWAS using a significance threshold of p = 0.0002 determined by a

Bonferroni correction of 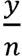, where y is alpha (α= 0.05) and *n* is the number of SNPs in the 15 priority loci (*n* = 219). Briefly, candidate genes were identified by querying the reference genome for coding sequences of genes within the priority loci. Annotated genes within these regions were compared using BLASTX against database of known genes and gene ontology information was catalogued. Special attention was paid to genes whose putative annotation was abiotic stress response. A more detailed version of the methods can be found in Supplementary Materials.

## Results

### Environmental variability across collection sites

Most of the 41 environmental variables showed variation across the sampled locations (Figure A in File S1). The Eastern Coast of Oaxaca and the Yucatan are significantly drier than the Western Coast of Oaxaca, though all three have relatively warm climates. The Central Valley receives less precipitation than the Western Coast, but it is significantly cooler than the coast and the Yucatan. Accessions from the Sierra Sur are located on the prevailing side of a rain shadow caused by the Sierra Madre del Sur Mountain range, so they receive about 150% the rainfall of accessions from the neighboring Eastern Coast and Central Valleys.

The PCA of the bioclimatic and edaphic variables (Figure 1) reinforces these differences; the first three principal components (PCs) explained 78.47% of the variance. The first PC (explaining 38.84% of variation) was largely associated with mean diurnal temperature range (bio2), annual temperature range (bio7), seasonality of precipitation (bio15), and the cation exchange capacity of the topsoil and subsoil. The second PC (explaining 21.73% of variation) was associated with annual mean temperature (bio1), the maximum temperature of the warmest month (bio5), the mean temperature of the warmest quarter (bio10), and the precipitation of the driest month (bio14)—variables of great interest in identifying local adaptation to drought stress. The third PC (explaining 17.89% of variation) is associated with annual precipitation (bio12), precipitation of the warmest quarter (bio18), pH of topsoil and subsoil, and the soil percentage of sand. A correlation analysis revealed that a number of the bioclimatic and edaphic variables were highly correlated (Figure B in File S1).

**Figure 1.**
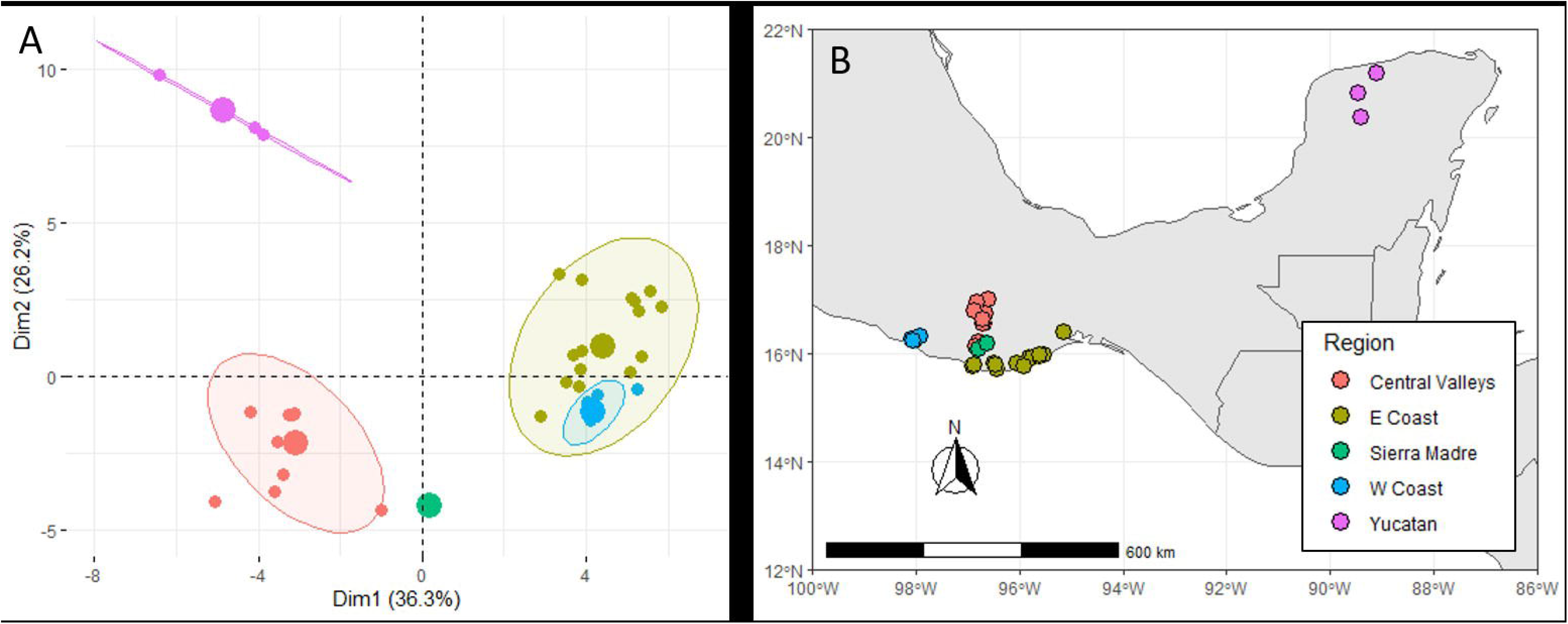
The five biogeographic ecozones from which we collected chile pepper accessions in Mexico. Identified on a principal component analysis (PCA) of elevation, bioclimatic variables, and edaphic variables (A), and by geographical distribution of accessions within those ecozones (B).

### F_ST_ and SPA Outliers

*F_ST_* and SPA outlier analyses were used to identify regions of the genome that may be under selection between populations and that demonstrate abrupt changes in allele frequency across the landscape, respectively, which may have contributed to local adaptation (Yang et al., 2012). Values for *F_ST_* range from 0 to 1; and SPA scores range from 0 to 13. Spatial distributions of genetic structure are often intertwined with population structure, yet only 34% of SPA and *F_ST_* outliers overlap with each other (Figure 2). However, the two measures are correlated (R=0.794, p <0.001) (Figures C & D in File S1) and identified 21 of the same loci (Figure 2).

**Figure 2.**
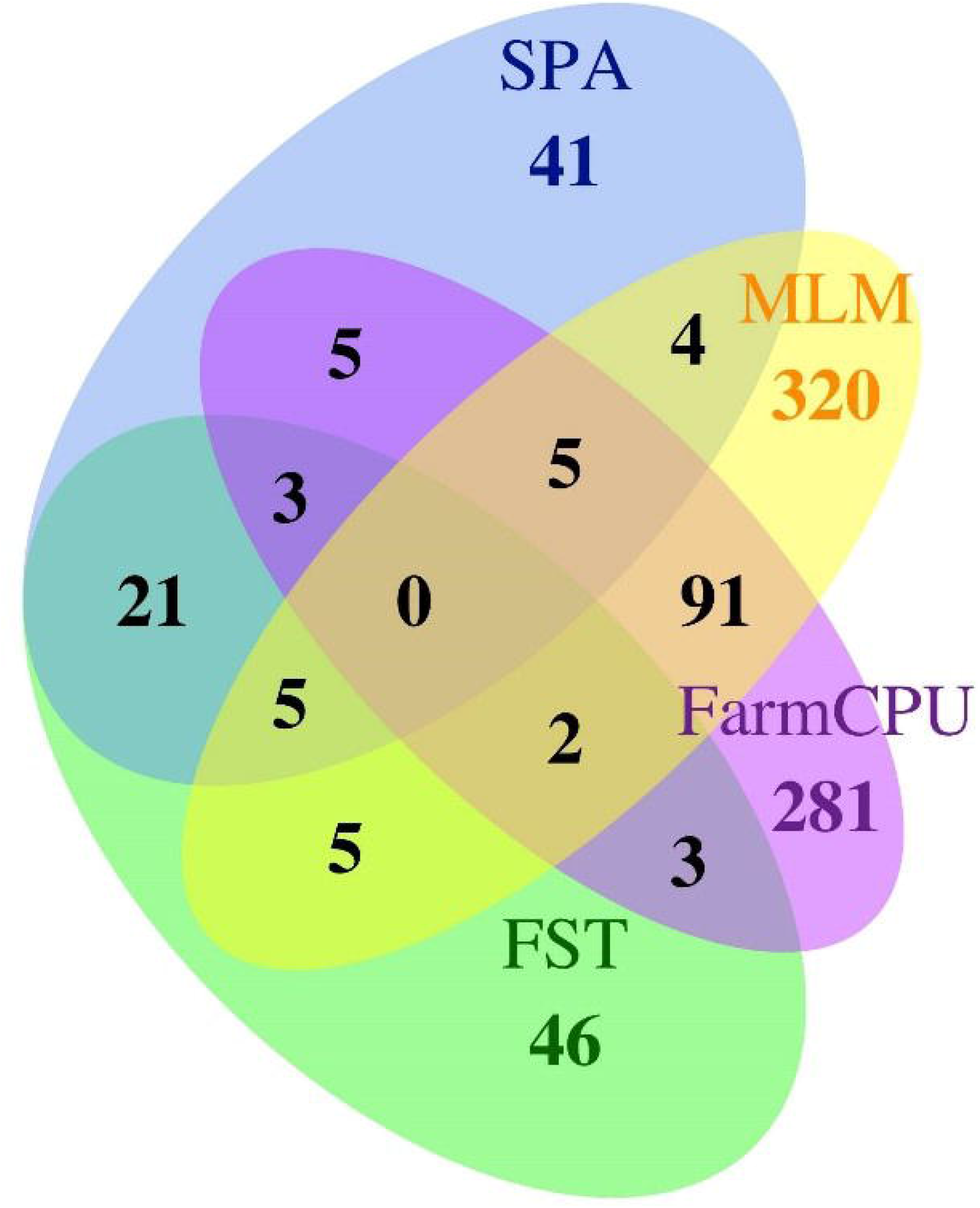
Summary of unique haplotype blocks identified using FS_T_ outliers (green), SPA outliers (blue), and two EAA models--MLM (yellow) and FarmCPU (purple). The overlap of the MLM and FarmCPU EAA, F_ST_, and SPA (red ellipse) were determined to be priority. Co-localization of loci occurred greater than expected by chance (P values < 0.00001; chi-square tests).

### Environmental association analysis

We will not present results from our EAA using the GLM and some analyses of some environmental variables using FarmCPU. Even with the inclusion of five principal components and the covariates of elevation, longitude, and latitude, confounding by population structure and relatedness was severe in the EAA performed using a GLM, producing highly inflated p-values (File S1). FarmCPU was unable to sufficiently account for population structure for analyses involving the following 12 (of 41) environmental variables: bio11, bio14, bio17, bio19, maximum DI, DI coefficient of variation, maximum PET, PET coefficient of variation, mean PET, subsoil organic matter content, subsoil cation exchange capacity and subsoil bulk density.

Nevertheless, the MLM (for all variables) and FarmCPU analyses (for the remaining 29 variables) controlled for these effects and detected significant associations with 1959 and 628 SNPs, respectively. These SNPs were within 432 and 390 linkage disequilibrium blocks (hereafter loci), respectively, for a total of 724 significant loci (Figure 2). Of the loci detected by MLM, 159 were related to temperature (bio1-11 and PET variables), 154 were related to precipitation (bio12-19 and DI variables), and 90 were related to soil variables, with significant overlap among these three groups (Figure E in File S1). Of the loci detected by FarmCPU, 179 were related to temperature, 30 were related to precipitation, and 149 were related to soil variables; as with the MLM, there was significant overlap among these three groups (Figure E in File S1).

### Identification of priority loci

Priority loci were determined to be those identified by two association models and one outlier analysis or two outlier analyses and one association model (Figure 2). Fifteen priority loci were identified on eight of twelve chromosomes (Table 2): one each on chromosomes three, four, five, and seven; two on chromosome nine; three on chromosome ten; and four on chromosome twelve. Two loci on chromosome 10 are just over 100kb from each other (339 677 bp to 470 407 bp and 573 263 bp to 742 206 bp).

**Table 1.**
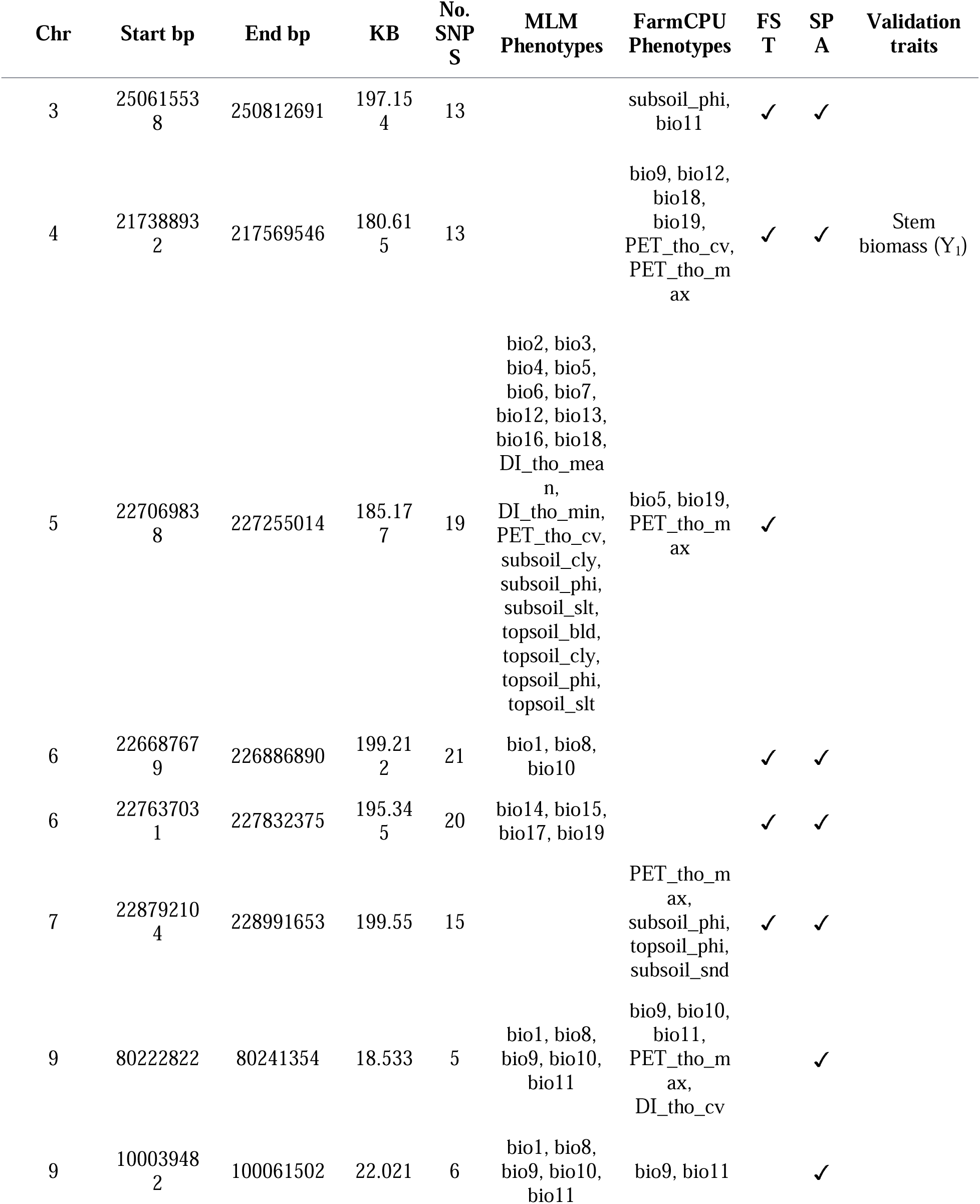
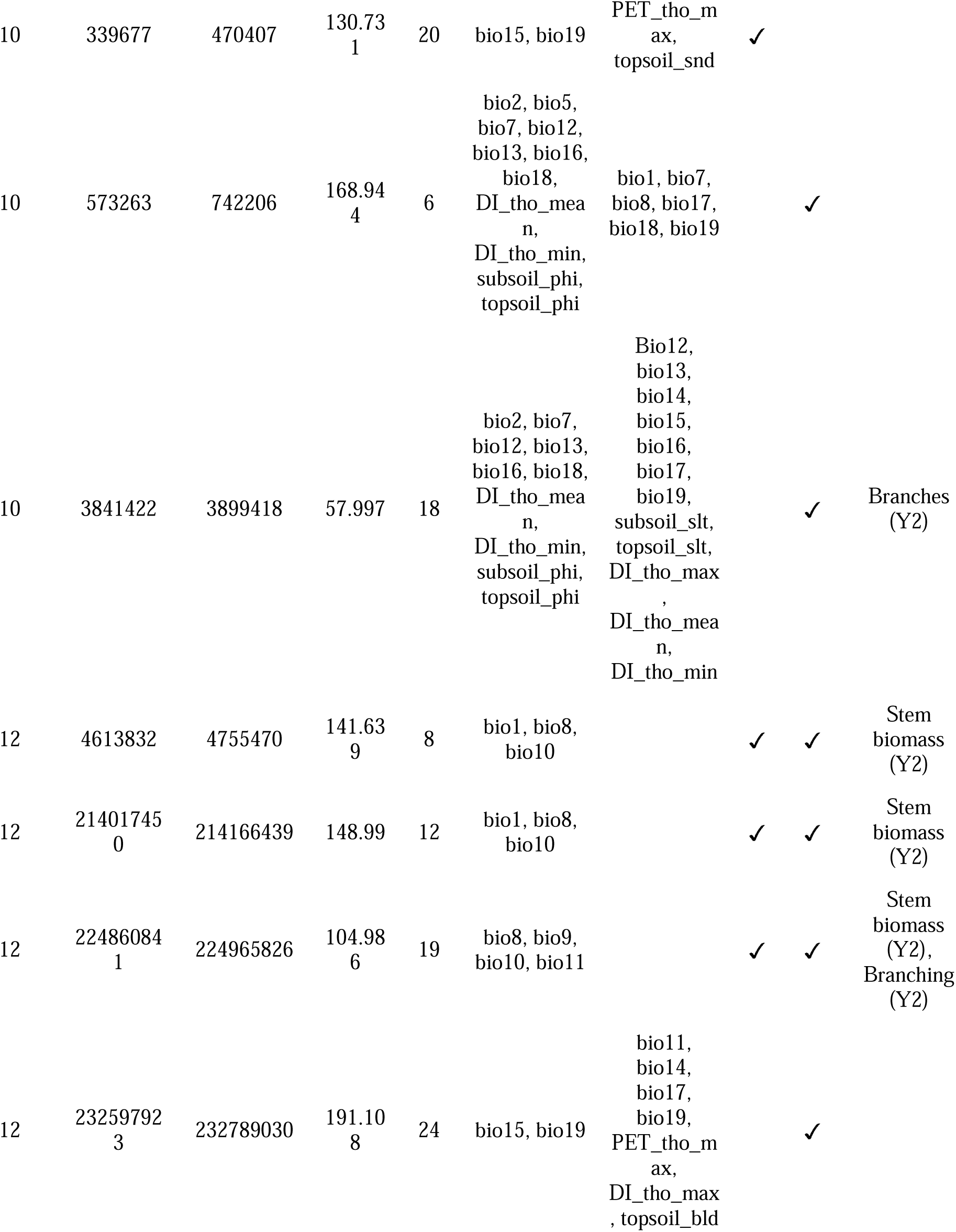
Fifteen priority loci as identified by the overlap between EAA, SPA, and FST.

**Table 2.** Loci estimated to explain more than 10% of the environmental variance across FarmCPU and MLM models.

| Model | Environmental phenotype | Block | SNP | +/- Allele | MAF | P-value | Effect | PVE |
| --- | --- | --- | --- | --- | --- | --- | --- | --- |
| FarmCPU | bio3 | 4_2854722 | S4_2854722 | G/A | 0.188 | $1.73 \times 10^{-07}$ | -0.36 | 11.6 |
| FarmCPU | topsoil_bld | 12_232597923 | S12_23266757<br>5 | G/A | 0.416 | $1.88 \times 10^{-13}$ | 0.474 | 10.4 |
| MLM | bio19 | 12_232597923 | S12_23266758<br>4 | A/G | 0.414 | $1.36 \times 10^{-07}$ | 0.577 | 17.3 |
| MLM | bio15 | 12_232597923 | S12_23266758<br>4 | A/G | 0.414 | $1.08 \times 10^{-07}$ | -0.576 | 14.4 |
| MLM | bio15 | 2_115018229 | S2_115156185 | T/A | 0.465 | $1.11 \times 10^{-06}$ | 0.608 | 10.3 |

Three priority loci identified in the overlap between the outliers of F_ST_, SPA, and the significant loci identified by the MLM GWAS model are located on chromosome 12 (Figure 3). They are primarily associated with temperature variables (e.g., annual mean temperature-bio1, mean temperature of the wettest quarter-bio8, and mean temperature of the warmest quarter-bio10).

**Figure 3.**
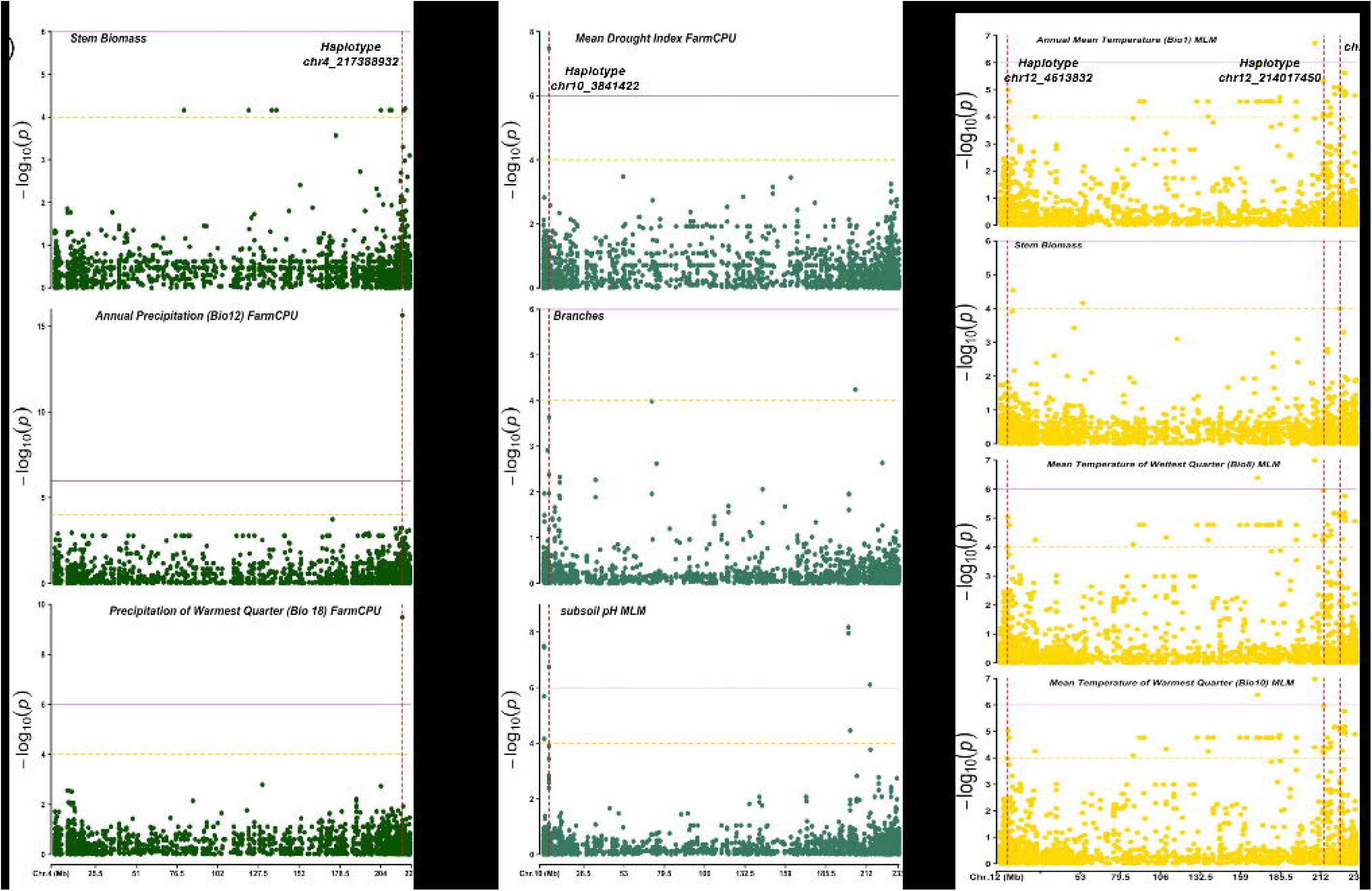
Examples of five validated loci (A) A validated locus on chromosome 4 which is also a SPA outlier. (B) A validated locus on chromosome 10 which is also a SPA outlier, (C) Three validated loci on chromosome 12.

Using the FarmCPU model, two loci were estimated to explain more than 10% of variance in the environmental phenotype (Table 2). One locus, located on chromosome 4, explained 11.6% of the variance of isothermality (bio3—calculated as the mean diurnal range divided by the annual temperature range). A second locus, on chromosome 12, was also identified as a priority locus in the overlap of EAA by MLM, SPA, and F_ST_. It explained 14.2% of the variance of topsoil bulk density. This locus was also identified by MLM and explained 17.3 and 14.4% percent of the variables precipitation seasonality (bio15) and precipitation of the coldest quarter (bio19), respectively. A second locus with a high estimated PVE in the MLM model was on chromosome 2. It explained 10.3% of the variance in precipitation seasonality.

### Association analysis of experimental phenotypes

We conducted a separate GWAS with three measures of stability calculated for five measured responses from the greenhouse based common-garden study: total biomass, fruit biomass, stem biomass, number of branching points, and fruiting (yes/no binary response at harvest). FarmCPU identified no significant associations across four haplotype blocks for the Y_2_ and Y_3_ stability measures for fruiting response, however both of these variables had poorly fitted QQ-plots and are not considered viable associations. The MLM model identified 27 significant associations across five haplotype blocks for the Y_2_ and Y_3_ stability measures of stem biomass production and number of branching points (Table 3). Furthermore, we evaluated genome-wide associations for drought tolerance at a more stringent threshold determined by 1000 permutations. Using the MLM and FarmCPU models, we found 29 and 21 significant SNP associations across 24 and 14 linkage blocks, respectively. These significant associations represent associations with three traits: stem biomass (Y_1_, Y_2_), fruit biomass (Y_2_), and total biomass (Y_2_). Significant associations were identified on all chromosomes except chromosome 2 and chromosome 9.

## Discussion

Using 32,623 high quality SNPs identified with GBS, we analyzed a collection of 190 georeferenced chile accessions of wild and cultivated landrace origin, from two states in southern Mexico. Our previous work confirmed genetic structure, genetic differentiation along an elevational gradient, and high genetic diversity among chile peppers in Oaxaca (Pérez□Martínez et al., 2022; Taitano et al., 2019). Here, using EAA, we identified significant genomic associations with bioclimatic variables, and, combined with measures of differentiation from population structure (*F_ST_*) and spatial differentiation (SPA), we identified 15 priority loci. Five of these priority loci were validated with experimental GWAS.

### Environmental association analysis and measures of genomic differentiation identify similar genomic regions

Combining results of different methods was expected to improve our ability to detect loci more likely to have been shaped by selection by local environmental conditions (Hoban et al., 2016; Sork et al., 2013). In our study, co-localization of 15 loci did not occur by chance alone (Figure 2). This significant level of co-localization validates each of the different analytical techniques applied. Yet, each technique also identified novel loci. For example, EAA methods identified 692 loci not identified by F_ST_ or SPA. The substantial divergence in results among methods highlighted the need to apply multiple approaches; thus, limiting the number of candidate loci to those most likely contributing to local adaptation (reviewed in Lasky et al., 2023; Rellstab et al., 2015)

### Managed ecosystems may not directly experience all environmental stressors

The geographic structure of the collections analyzed was limited to three distinct transects: one that follows the increasing precipitation gradient going east to west along the Oaxacan coast, one which parallels the decreasing temperature and increasing elevation from the Oaxacan coast to the Central Valleys, and one along the coast of the Yucatan. These three transects cover a relatively small geographic area, and less than half of *Capsicum*’s climate envelope across Mexico (Khoury et al., 2020). As pointed out by (Anderson et al., 2016), environmental association analyses are dependent on quality, high resolution global datasets to reduce “phenotyping error.” As bioclimatic variables are treated as phenotypes, we work under the assumption that these variables represent selective pressures on the accessions studied. This connection is rather straightforward in wild and feral populations, but most of the populations studied here come from managed ecosystems (though the degree of management varies). In managed systems, farmers may buffer some environmental stressors (e.g., through irrigation or greenhouses and other controlled environment systems). However, there is still likely selection (both natural and human-mediated) from year to year for plants that perform the best in the target environment, contributing to local adaptation of the farmer’s landrace populations (Bandillo et al., 2015; Lasky et al., 2015; Russell et al., 2016).

### Five loci were phenotypically validated

Using the approach of prioritization by co-localization of loci, identified by multiple landscape genomics methods, led to the validation of 33% (5/15) of the prioritized loci. Attempts to phenotypically validate loci identified by landscape genomics has been rare in crop plants (e.g. Lasky et al., 2015; Russell et al., 2016). Through phenotypic GWAS, we were able to validate five of the fifteen priority loci identified by the co-localization of EAA, SPA, and F_ST_. Of these five validated loci, three appeared to be related to temperature in their environment of origin, one to water and one to both. Those related to temperature were all found on chromosome 12. The locus related to water was found on chromosome 4 and showed a large phenotypic response in stem biomass under water deficit. On chromosome 10, the locus was related to water, temperature, and silt content of the soil, as well as branching under water deficit. We may be seeing the genetic signature of multiple abiotic stress adaptations in these accessions, which could be linked to different phenotypic syndromes (e.g., plant architecture; McCoy et al. 2022). Genomic regions associated with multiple environmental variables may be indicative of pleiotropic or coincident adaptation to varied abiotic stresses (R Wang et al., 2023). Further investigation of drought responses can lead to an understanding of the genetic architecture and phenotypic networks that control this complex trait.

### Priority loci reveal unique genetic variation

The QTL identified on chromosome 4 (bp 217 388 932 to 217 569 546) was particularly interesting as it was significant in multiple EAA analyses, FST, and SPA. Additionally, it was phenotypically validated with significant association with stem biomass under water deficit conditions. The haplotype associated with low rainfall but also low temperature and low PET was identified solely from *Chile de Agua* in the ‘Central Valleys.’ Among the priority loci identified in this study, there was often a distinct Central Valleys haplotype (e.g., group B in Figure 4). The ‘Central Valleys ‘accessions had a distinctive architecture with elongated and thick herbaceous stems indicating a maladaptation to water deficit, as expected by its low PET origin. As discussed in Taitano et al. (2019), *Chile de Agua* had the lowest average nucleotide diversity (π) of the major landraces included in their collections, which were also used in this study. A full 64 of our 190 studied lines were of the landrace *Chile de Agua* from the Central Valleys. Taken together, the high degree of homogeneity within the *Chile de Agua* accessions and the high number of plants with the same haplotype, contributes to a fixed haplotype across the geographic region, showing the need for increased conservation and the utility of these collections to identify novel variations related to abiotic stress tolerance. Here we see there is a such a need for validation as looking at environmental traits alone (i.e., rainfall and PET) provide a shadowy picture that is only illuminated in the presence of a phenotypic screen of promising material.

**Figure 4.**
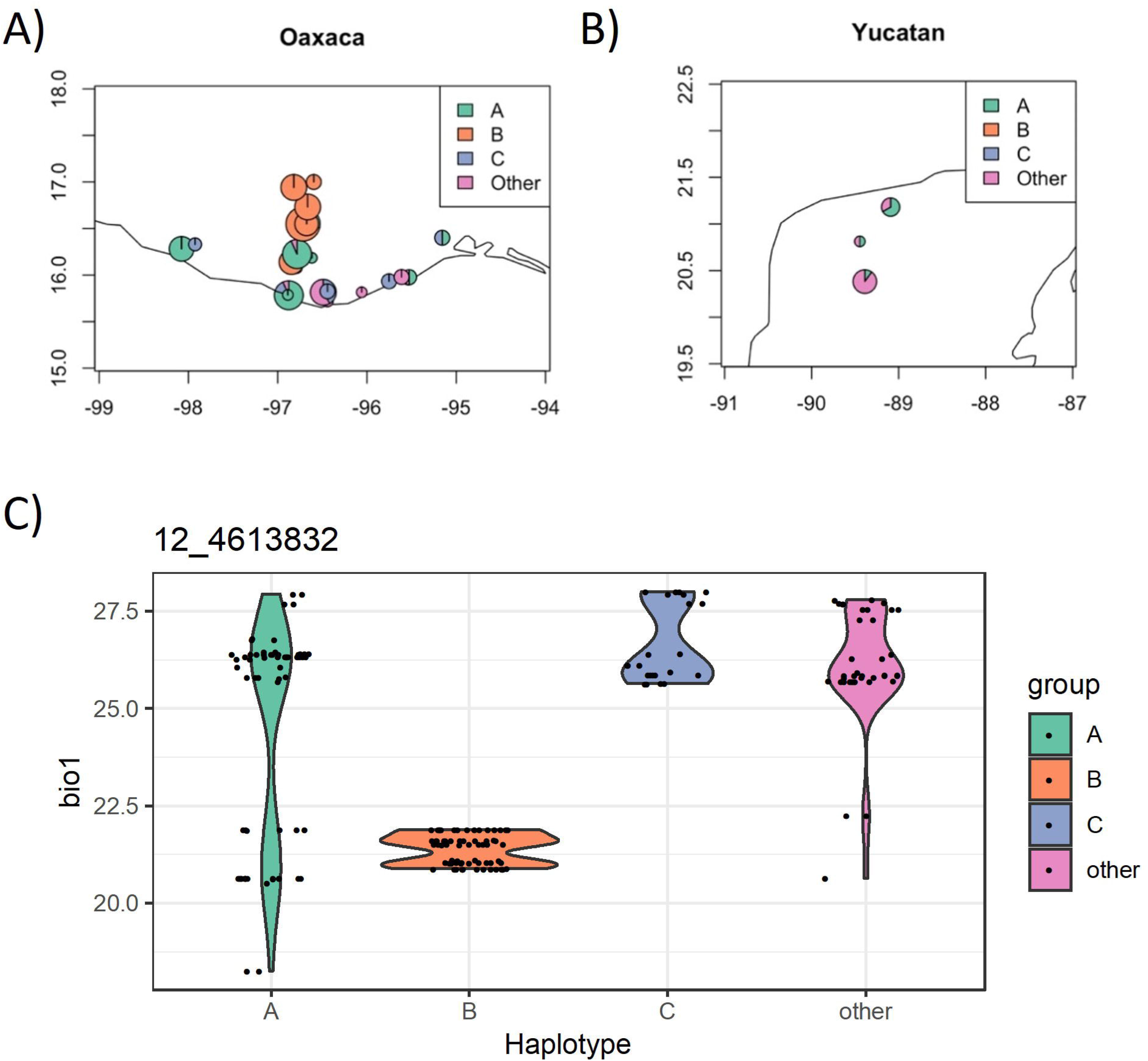
EAA identified between basepairs 4 613 832 and 4 755 470 on chromosome 12 of the *Capsicum* genome. (A) The MLM model identified significant associations at this locus for annual mean temperature (bio1), mean temperature of the wettest quarter (bio8), and mean temperature of the warmest quarter (bio10). (B) The FarmCPU model identified a significant association for mean temperature of the wettest quarter (bio8). Geographic location. Of individuals with Haplotype A (teal), Haplotype B (orange), Haplotype C (lavender), and other haplotypes (pink) across Oaxaca (C) and Yucatan (D). Violin plot of haplotype distribution for annual mean temperature (bio1, E), mean temperature of the wettest quarter (bio8, F), and mean temperature of the warmest quarter (bio10, G). The units for the y-axis of all violin plots is mm.

### Candidate genes within key haplotype blocks have putative roles in abiotic stress response

Genes within haplotype blocks defined by the 15 priority loci are possible candidate genes involved in environmental adaptation and abiotic stress tolerance (Table S2). Here we focus our discussion on the five validated priority loci. Candidate genes were identified for four of the five validated priority loci based on physical position and predicted function. Within the 28 genes predicted to fall within the associated genomic sequence of the five validated priority loci, there were six candidate genes with predicted biological functions related to stress response.

On Chromosome 4 (217,388,932 to 217,569,546 bp), three candidate genes were identified. These included two genes (*CA04g20990* and *CA04g21000*) predicted to encode jasmonic acid carboxyl methyltransferases (JMTs). *JMT* genes act to carboxylate jasmonic acid to form methyl jasmonate, which is known to induce defense response genes. *JMT* genes, specifically, have been well studied for their induction following wounding and herbivory (Song et al., 2005). Further, this group of genes has been related to leaf senescence and the hypersensitive response, which may act in response to stress (Huang et al., 2020, p. 2; Raffaele et al., 2008) The third gene (*CA04g21020*) is predicted to encode an AAA-family ATPase, a member of a large family of proteins known for their diverse cellular functions (Confalonieri & Duguet, 1995). *CA04g21020* has homology with a gene predicted to encode a mitochondrial BCS1-like protein specifically upregulated in salt-stressed roots in Arabidopsis (Ma et al., 2006).

On chromosome 10 (3,841,422 bp to 3,899,418 bp), within the validated locus associated with temperature stress and drought stress—among other environmental variables, the gene *CA10g01780* is predicted to encode a protein putatively involved in UV-damage excision repair (Soderlund et al., 2009). Indeed, previous work has shown that with domestication, the cultivation of chile peppers has moved to higher elevations than wild *C. annuum* populations (>1500 meters above sea level) (Martínez-Ainsworth et al., 2022).

On Chromosome 12 (4,613,832 bp to 4,755,470 bp), the validated locus associated with temperature includes *CA12g02160,* predicted to encode an aspartyl protease and shown in Arabidopsis to respond to oxidative stress (Stanley Kim et al., 2004). An increase in the amount of reactive oxygen species (i.e., oxidative stress) is a well-known response to drought and increased temperature (Hasanuzzaman et al., 2013). Having the ability to deal with these compounds provides an additional layer of defense (Qu et al., 2013).

A second validated priority locus on Chromosome 12 (214,017,450 bp to 214,166,439 bp), which was associated with environmental variables related to temperature, encompasses *CA12g16320* predicted to encode a member of the canonical disease resistance protein family, nucleotide binding-leucine rich repeat proteins (NLRs). While this large protein family is well studied for its role in biotic stress response (reviewed in McHale et al., 2006), there is also significant crosstalk and integration among biotic and abiotic stress signal transduction (reviewed in Sijo and Loo, 2020). For example, in Arabidopsis, through crosstalk between the salicylic acid and abscisic acid pathways, the expression of the NLR, *Activated disease resistance 1*, enhances resistance and drought tolerance at the expense of salt and heat tolerance (Chini et al., 2004).

## Conclusions

Identification of genomic loci and alleles associated with environmental adaptation in wild and landrace accessions collected across their center of crop origin can inform strategies of how-to best conserve and leverage genetic resources for climate resilience. Here, we identified and phenotypically validated QTL associated with multiple environmental variables, with particular focus on variables related to abiotic stress tolerance, especially drought tolerance. We explored a unique collection which represents a geographic gradient of landraces from the center of origin of the species. Analyses like this one can help shed light on possible paths forward for conservation and use of public data and resources as well as providing insight into adaptation in plants.

## Data Availability

To protect the rights of the indigenous peoples who have cultivated and cared for these crops, germplasm and coordinates for the original collections used in this study are not available for distribution. However, the haplotype sequences of the priority loci have been provided for comparing the representation of these loci in other germplasm samples. Input data and scripts for the analyses are being uploaded to Dryad (doi in progress).

## Acknowledgements

This project was funded by a SEEDS Graduate Research Grant provided by the Ohio Agriculture Research and Development Center (OARDC) (2015100) and a SEEDS OARDC Research Enhancement Grant (2016056). We acknowledge the Center for Applied Plant Sciences and Ohio State University for funding the collection trips. We would like to thank the USDA-AFRI, Physiology of Agricultural Plants section for support under grant number 2017-06351, “Genetic structure and mechanisms of drought adaptation in Capsicum”. L.J.B. thanks the support of the Programa de Apoyos para la Superación del Personal Académico (UNAM-DGAPA) for the financial support for the sabbatical stay at The Ohio State University during which this manuscript was finished and revised. This work was supported in part by the U.S. Department of Agriculture, Agricultural Research Service.

## Notes

### Competing Interest Statement

The authors have declared no competing interest.

### Summary of Updates

Data availability statement updated to indicate current status of data archiving. Supplemental file 2 was updated with accurate table names.

