## Supplemental File 1 for "Genomic signatures of adaptation to abiotic stress from a geographically diverse collection of chile peppers (Capsicum spp.) from Mexico"

### Supplementary Material

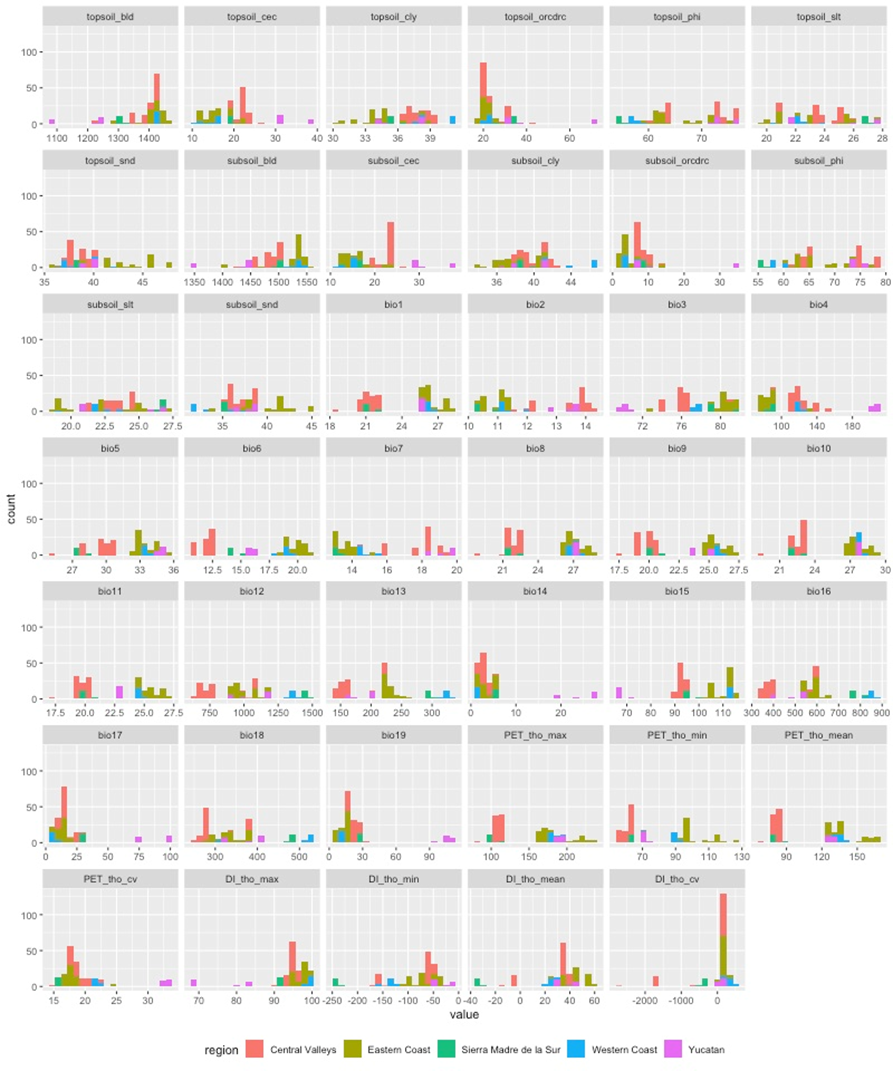
Figure A. Distribution of environmental phenotypes by ecozone. Colors represent ecozone of origin.

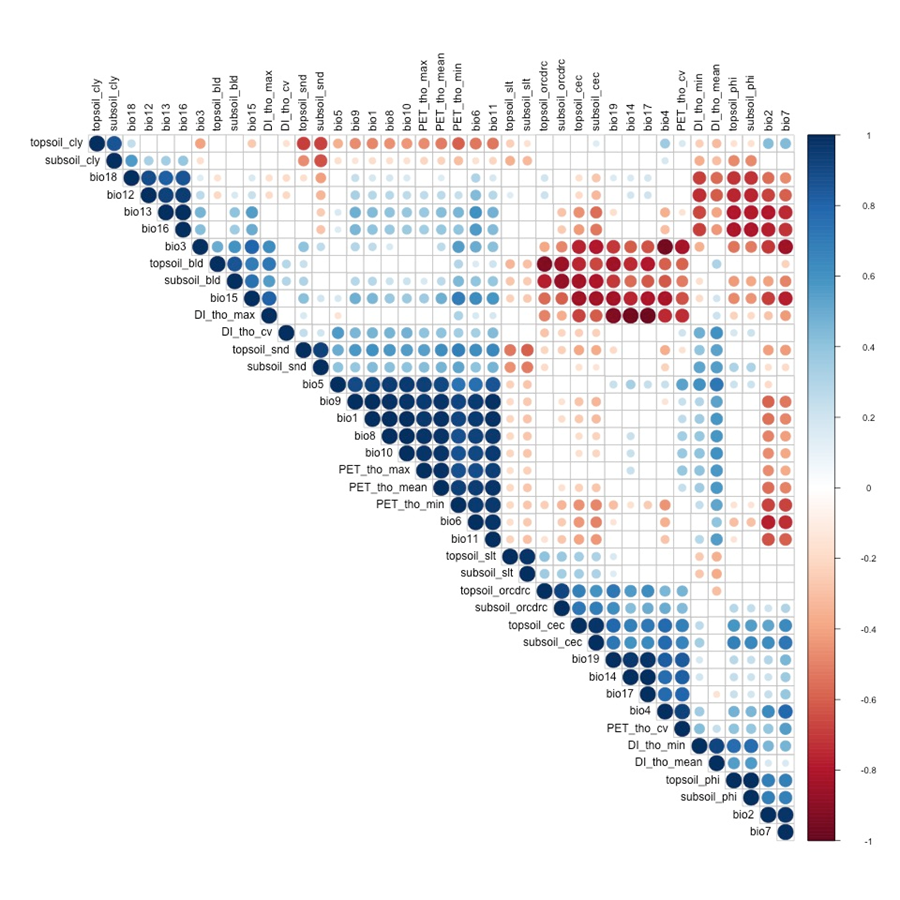

Figure B. Pearson correlation plot of all environmental phenotypes. Size and color of the circles relates to the correlation strength of significant correlations. Correlations where p>0.05 are blank.

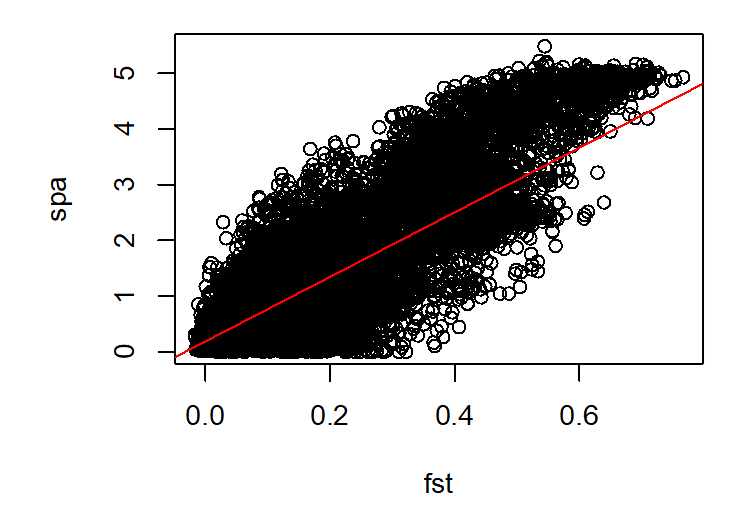

Figure C. Plot of SPA scores against F_ST_ values (R = 0.794, p <0.001).

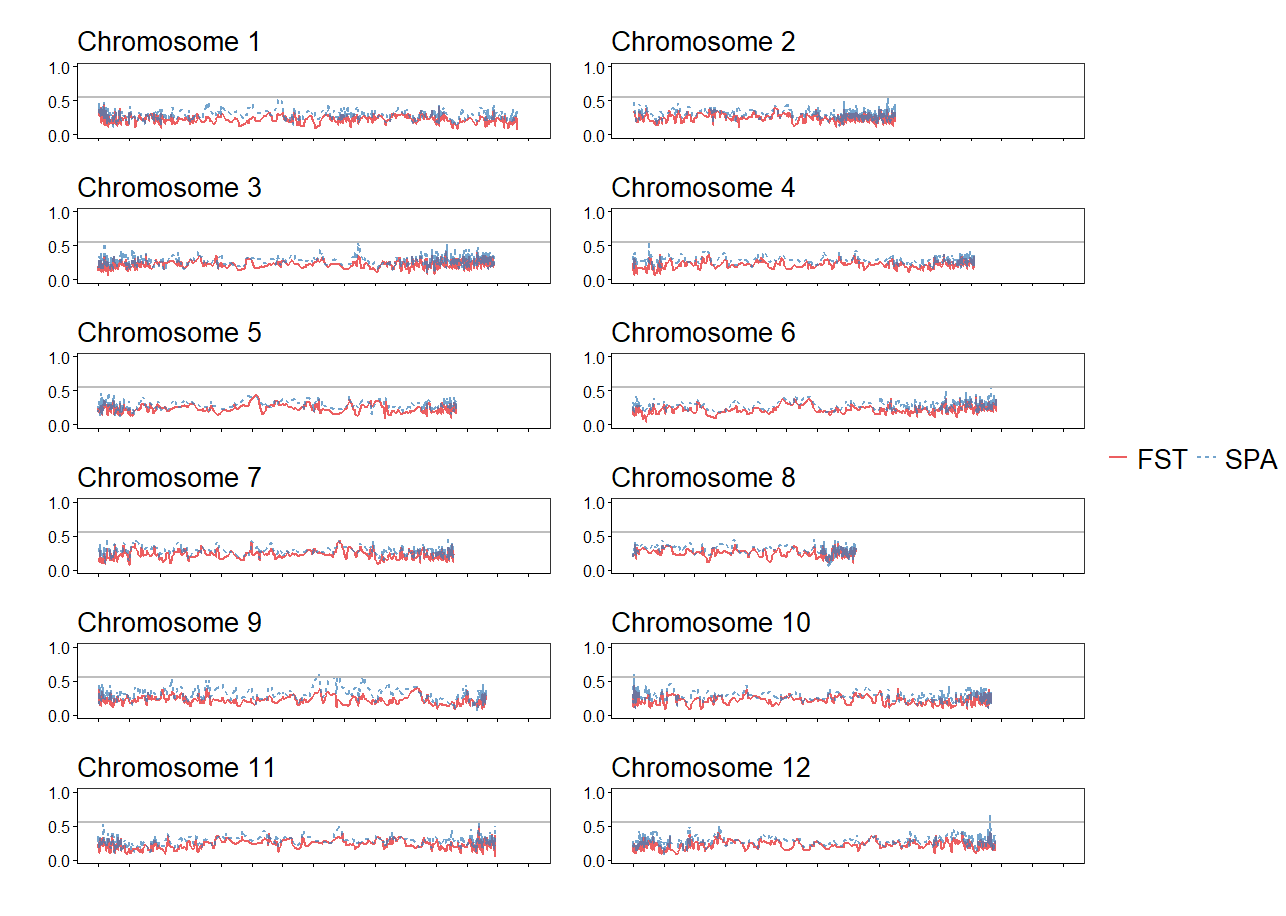

Figure D. Sliding window of SPA index (SPA / max SPA value, blue) and FST (red) across the genome. A threshold of 0.55 identifies the top SPA outliers.

A) MLM
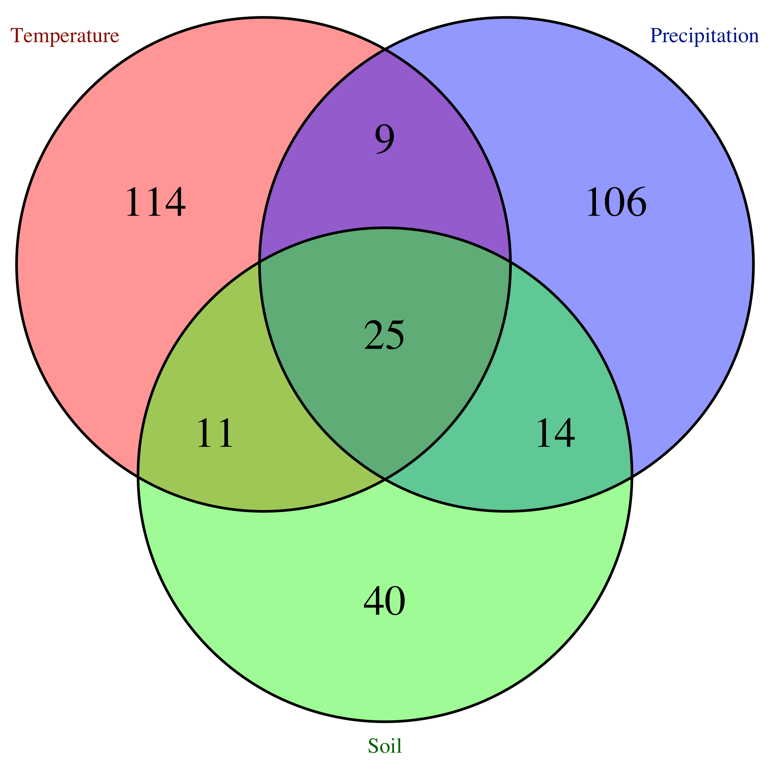

B) FarmCPU
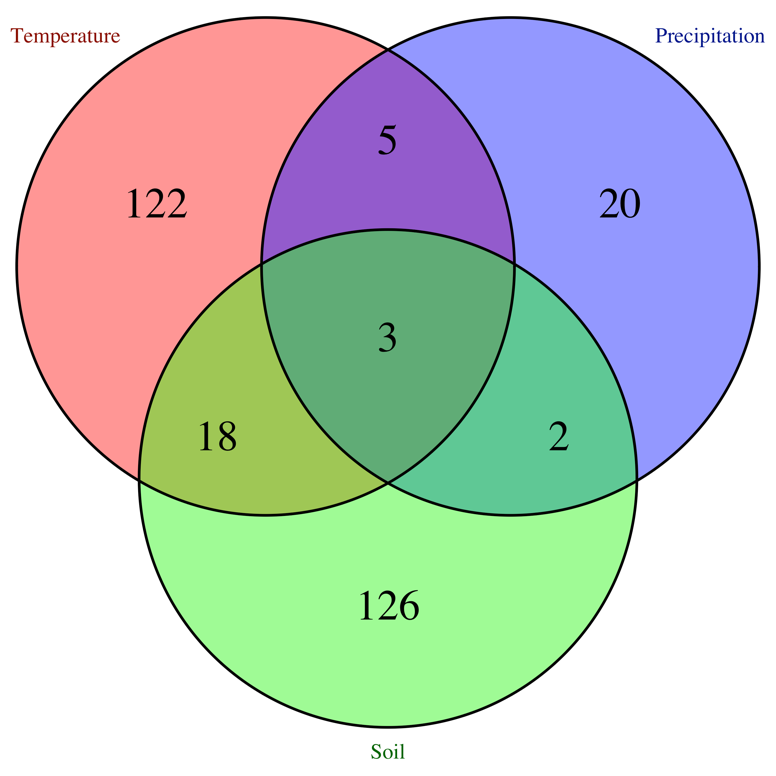

Figure E. Number of loci identified for each type of environmental data point for the MLM model (A) and FarmCPU model (B). Temperature variables (red) include bio1-bio11 and the calculated PET measures; precipitation variables (blue) include bio12-bio19 and the drought index variables; soil variables (green) include the ISRIC data points for topsoil and subsoil.

Complete Materials and Methods
*Plant material and genetic data*
In 2013, we collected pepper accessions in two states of Mexico: throughout Oaxaca and in northern Yucatan (Figure 1). In Oaxaca, we first collected at 13 sites along a north-south transect from the southern tip of the coastline in Pochutla, increasing in elevation to the Central Valleys near Oaxaca City. Our second Oaxacan transect was oriented east to west along the coast, included 12 collection sites, and followed a precipitation gradient (increasing towards the west). In addition to crossing climatic gradients, the Oaxacan transects spanned ethnic and language groups. Collections were also made from three villages in the Yucatan (Table 1).
DNA was extracted, genotyped and processed as described for the “Mexican Collection” in Taitano et al. (2018). Briefly, 50 mg of young leaf tissue was collected from adult chile pepper plants, flash frozen in liquid nitrogen, lyophilized and ground to a fine powder using a Geno/Grinder 2000 (SPEX, Metuchen, New Jersey, USA). DNA extraction was performed using QIAGEN’s DNEasy 96 Plant Kit® (Valencia, California, USA), following the manufacturer’s recommendation. DNA was eluted into 100 μl TE pH 8. Genotyping-by-sequencing (GBS) libraries were constructed for each genotype according to the Elshire method and 48 libraries were pooled. Two pools were sequenced on the NextSeq platform and two additional pools were sequenced on the HiSeq 2500 platform. The TASSEL GBS Pipeline 5.2.3 (Glaubitz et al., 2014) was used to call single-nucleotide polymorphisms (SNPs) from Illumina sequence data. The *C. annuum* cv. CM334 reference genome was used for read alignment with Bowtie2 (Langmead and Salzberg, 2012); a minor allele count of three reads per minor SNP allele was required to call a SNP.
*Bioclimatic and edaphic variables*
We queried two global datasets: WorldClim2 (Fick and Hijmans, 2017) and ISRIC (World Soil Information database; Hengl et al., 2017) for climate and edaphic data, respectively, using the latitude and longitude coordinates recorded with the original collections. WorldClim2 was queried for monthly estimates of precipitation, wind speed, solar radiation, water vapor pressure, and maximum, minimum, and average temperature at a resolution of 30 arc seconds (approximately 1-km^2^ at the equator). Nineteen summary bioclimatic variables describing monthly, seasonal, and yearly variation in temperature and precipitation were also accessed at 30 arc second resolution. ISRIC was queried for seven edaphic variables at a resolution of 1-km^2^: bulk density, cation exchange capacity (CEC) (cmolc/kg), organic carbon content (fine earth fraction) in permilles, soil pH ([pH] × 10 in H_2_O), percent sand, percent silt, and percent clay. We calculated bulk estimates for topsoil (0-30 cm) and subsoil (30-200cm) from measurements at seven depths using weighted averages (Hengl et al., 2017; per methodology in Anderson et al., 2016; Bandillo et al., 2017).
We used the Thornthwaite approximation to estimate monthly potential evapotranspiration (PET) at each germplasm collection site Thornthwaite and Mather, 1955; Thornthwaite et al., 1957). These calculations were made in R version 3.3.3 (R Core Team, 2017) using the package ‘SPEI’ (Beguería and Vicente-Serrano, 2017) with climate data from WorldClim2. We then calculated a drought index (DI) as follows using precipitation (P) data also from WorldClim2.
${DI}_{j,i}=100\left( \frac{PET-P}{PET} \right)$
The maximum, minimum, mean, and coefficient of variance of monthly PET and DI estimates were calculated for each site.
The 19 bioclimatic variables, topsoil and subsoil data, and PET and DI values were scaled to a mean of zero and a standard deviation of one since the genome-wide association analysis assumes a normal distribution of traits. A principle components analysis (PCA) was conducted using the package “FactoMineR” (Lê et al., 2008) and visualized using the “factoextra” (Kassambara, 2018). Pearson correlation coefficients were calculated in R and visualized with the package “corrplot” (Wei and Simko, 2017). Histograms for each variable were created to examine the distribution (Figure A in File S1).

Statistical Analysis

*F_ST_ and SPA outlier analyses*

A F_ST_ outlier scan was used to identify loci with differential allele frequency across groups. We used six genetic clusters (i.e., k = 6) as identified in a fastSTRUCTURE analysis (Raj et al., 2014; Taitano et al., 2019). Theta (θ), the variance-based *F_ST_* estimate of Weir and Cockerham (1984), was estimated using the R package ‘hierfstat’ (Goudet, 2005). For visualization, *F_ST_* was averaged in sliding windows, with a window size of 10 and a step of three SNPs. The 100 SNPs with the highest score (above 99.7 percentile) were identified as outliers.
Spatial ancestry analysis (SPA) was conducted to detect loci showing steep gradients in allele frequency across space (Yang et al. 2012). SPA assesses samples across continuous geographic and environmental space and estimates an allele frequency function that is projected onto geography. Loci with steep allele frequency gradients are likely the products of selection (Yang et al. 2012). SNPs with SPA scores above the 99.9th percentile were identified as outliers with steep enough allele frequency gradients to have been the subject of selection. SPA comparisons are useful when individual relationships are driven by isolation by distance, as the gradient function incorporates geographic and genetic gradients to identify local adaptation clines (Yang et al. 2012). Non-biallelic SNPs were filtered out for a total of 29,986 SNPs. A supervised mapping analysis with known location was conducted. The top 100 scores (above 99.6 percentile) were identified as outliers.
*Environmental association analysis*
Genome-wide association studies (GWAS) use statistical models to identify associations between a genetic marker and a phenotype. There are a number of statistical models that attempt to control false-positives (type I error) in GWAS by incorporating variations of kinship matrices and principal components to account for population structure. The work presented here was conducted using the ‘MVP’ package in R, which combines the models GLM (general linear model), MLM (mixed linear model), and FarmCPU in a memory-efficient, visualization-enhanced, and parallel-accelerated tool (Yin et al., 2017). The GLM model (also known as the Q-model) incorporated principal components (PCs) derived from genetic markers (in this case, SNPs)in a fixed model to account for population structure (Price et al., 2006). The MLM is more computationally intensive and accounts for both population structure (using PCs as in the GLM) and kinship between individuals (Price et al., 2006; Yu et al., 2006). To reduce the computational burden of the complex MLM model, a variety of algorithms have been developed; we used the Efficient Mixed-Model Association (EMMA) (Kang et al., 2008; Zhang et al., 2010). FarmCPU uses a multi-locus mixed model (MLMM) which incorporates multiple markers simultaneously as covariates in a stepwise MLM. This partially removes the confounding between testing markers and kinship. It helps control for the effects of large effect loci so that underlying smaller-effect loci can be identified (Kang et al., 2008; Zhang et al., 2010). As GWAS models are not designed to account for spatial structure between individuals, covariates of elevation, latitude, and longitude for collection sites were included in all models.
The haplotype block of the GWAS population was estimated in PLINK v1.9 (Chang et al., 2015) with the following settings: ‘–no-parents –allow-no-sex –blocks’. PLINK uses the default procedure of Haploview, which ignores markers with minor allele frequency (MAF) < 0.05. In Haploview, 95% confidence bounds on *D′* are generated, and each comparison is called "strong LD (linkage disequilibrium)", "inconclusive" or "strong recombination". *D′* is the normalized coefficient of linkage disequilibrium (*D*), which represents the difference between the frequency of a haplotype and its probability. A block is created if 95% of informative (i.e. non-inconclusive) comparisons are "strong LD". By default, all possible blocks are sorted largest to smallest, and non-overlapping blocks are added in order of decreasing size (Barrett et al., 2005). Haplotypes represented by more than 10% of individuals were visualized for each priority loci.
Since the environmental phenotypes used in the EAA are not normally distributed, a significance threshold was estimated for each variable using 1000 permutations in the MVP package. This threshold ranged from 5.45 × 10^-12^ to 3.26 × 10^-5^. The permutation test breaks the relationship between genotypes and phenotypes with each permutation, and the output is a vector of the minimum p-value for each permutation. The 95% quantile value of the vector is recommended as the threshold for identifying significant markers in the output of MVP models. For each association, the SNP with the lowest p-value in each haplotype block was used to estimate the percent variance explained (PVE). PVE was estimated for each locus as the difference between the R^2^ of a linear model including all significant loci and the R^2^ of a linear model without that loci. In order to test full linear models, without filtering significant SNPs due to missing data, PVE was estimated using a LD-KNNi imputed dataset calculated in Tassel (v5.0) (Bradbury et al., 2007; Glaubitz et al., 2014).
*Phenotypic GWAS*
A greenhouse-based common garden study with manipulated well-watered and water deficit treatments provided phenotypic trait data for our phenotypic GWAS. Accessions were grown in a greenhouse on the campus of the Ohio State University in Columbus, Ohio, USA. The greenhouse was maintained at 29.5˚C (85˚F) with 12/12 hours light/dark. We used a randomized complete block design (RCBD) with three replicates in time. A fourth replicate had to be aborted due to viral infection, so fewer replications than planned may have limited our statistical power to determine some differences across landraces and cultivation systems. Seeds were started in flats and transplanted to six-liter pots after six to eight weeks. Transplant time was adjusted to account for seedling size—despite supplemental light, this varied due to seasonal light intensity. Thirty-six grams of Osmocote^®^ Plus slow-release (3-4 month) fertilizer was incorporated into each pot (The Scotts Company, Marysville, OH). This fertilizer has a NPK analysis of 15-9-12, and it provides micronutrients magnesium, sulfur, boron, copper, iron, manganese, molybdenum and zinc. All pots were watered well for one week after transplant, then a drought treatment was applied using pressure-compensating emitters. Pots under the water-limited treatment always received 1/3 of the water supplied to the well-watered treatment. The irrigation time was adjusted throughout each block to maintain pots in the well-watered treatment at pot capacity without leaching. Plants in the third replication required longer to develop because of the lower light intensity in winter months. The greenhouse was sprayed regularly with insecticides to control thrips.
As the size of each block did not permit all measurements to be completed by one person, or in a single day, measurements taken on separate days or by more than one person were collected so as to be confounded with the random factor “bench”, which otherwise accounts for environmental variation across the greenhouse within each block.
The area of the upper-most, fully expanded leaf on each plant (destructive) was measured at two points during each block using a table-top leaf area meter (LI-3100, LICOR Inc, Lincoln, Nebraska, USA). Leaves were dried in a biomass oven, allowed to equilibrate to ambient conditions, and weighed to calculate specific leaf area (SLA), the ratio of leaf area to dry mass. At the end of each block, plants were scored on a scale of one to five for the severity of aborted leaves, with one representing no dropped leaves and five representing 50% or more leaves aborted. The number of fruit was recorded (for two of three blocks), and all above-ground biomass was harvested and dried in a biomass oven. The dry biomass was partitioned into stems, leaves, and fruit. The height of the dry plant and number of branching nodes for each plant was also recorded.
Aside from fruiting data (fruit number, fruit mass, fruit index), all data was normally distributed. To account for the Poisson distribution of the fruiting data, a binary response of fruiting (1) or not fruiting (0) at harvest was tested using a generalized linear mixed model with a binomial link function. Fruit number, fruit mass, and fruit index data for plants that had fruited by harvest (i.e., those that had non-zero values fruiting) were then analyzed without transformation using the mixed model above.
GWAS using those phenotypic data can provide additional priority loci that can be compared to those identified from *F_ST_*, SPA, and EAA. We analyzed a linear mixed model using restricted maximum likelihood with the R packages lmerTest (version 3.0) (Kuznetsova et al., 2016), emmeans (version 1.2) (Lenth, 2018), and their dependencies: lme4 (version 1.1) (Bates et al., 2015), and matrix (version 1.2) (Bates and Maechler, 2017). The model tested the effect of line, the water treatment, and their interaction as fixed factors. As random factors, we included blocks over time and bench nested within block. Bench nested within block accounted for variation across the greenhouse during each temporal block. BLUPs (best linear unbiased predictors) of total biomass, stem biomass, fruit biomass, branching, and fruit number were calculated for the 156 lines under each water treatment and combined in three functions of trait stability under the drought treatment:
1) $Y_{1}=\mu_{Control}-\mu_{Drought}$
2) $Y_{2}=\left[ \mu_{Control}-\mu_{Drought} \right]/\mu_{Control}$
BLUPs could not be estimated for leaf biomass or fruit number. The experimental phenotypes were analyzed in a GWAS using the same settings as described above for the environmental association. We used a significance threshold of p = 0.0002 determined by a Bonferroni correction of y/n, where y is alpha (α = 0.05) and *n* is the number of SNPs in the 15 priority loci (*n* = 219).
We calculated broad-sense heritability (*H^2^*) for each trait on a landrace basis using the following equation:
$H^{2}=\frac{V_{G}}{\left( V_{G}+V_{GTt}+V_{er} \right)}$
where *t* is the number of treatments, *r* is the number of replications, and, in a random-effects model, *V_G_* is the variance due to genotype (in this case, population), *V_GT_* is the variance due to the G×E interaction (in this case, population × water treatment), and *V_e_* is the residual variance.
