## Supplemental File 3 for "Genomic signatures of adaptation to abiotic stress from a geographically diverse collection of chile peppers (Capsicum spp.) from Mexico"

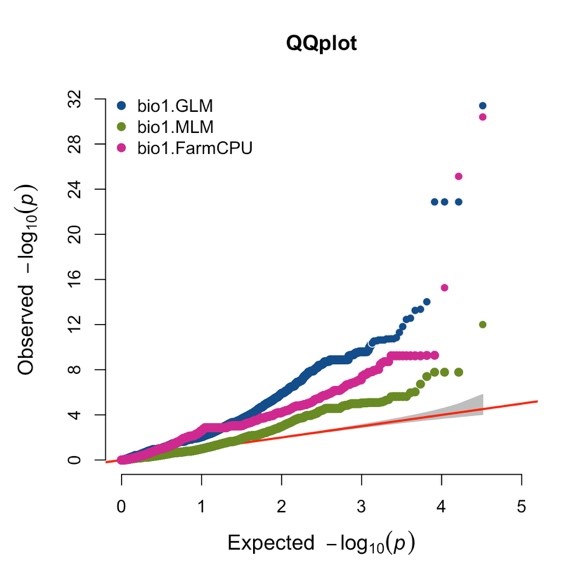

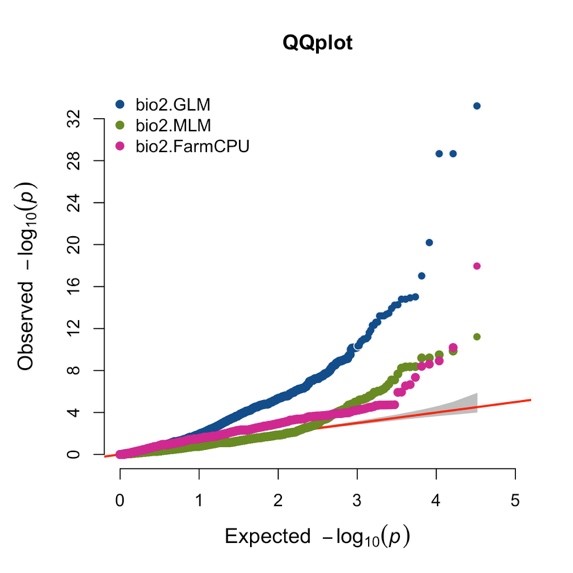

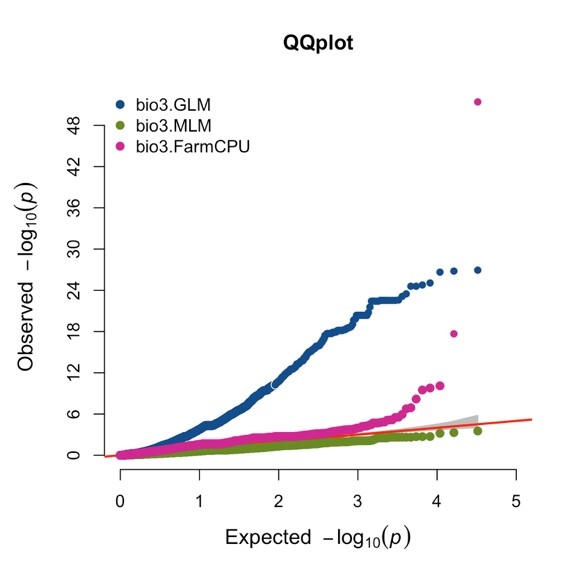

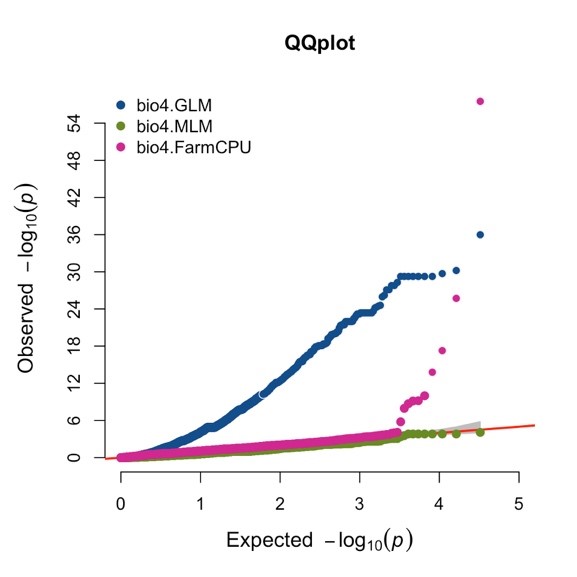

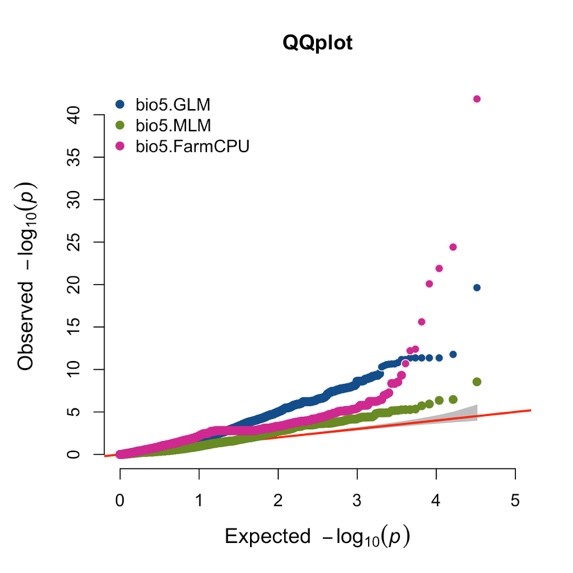

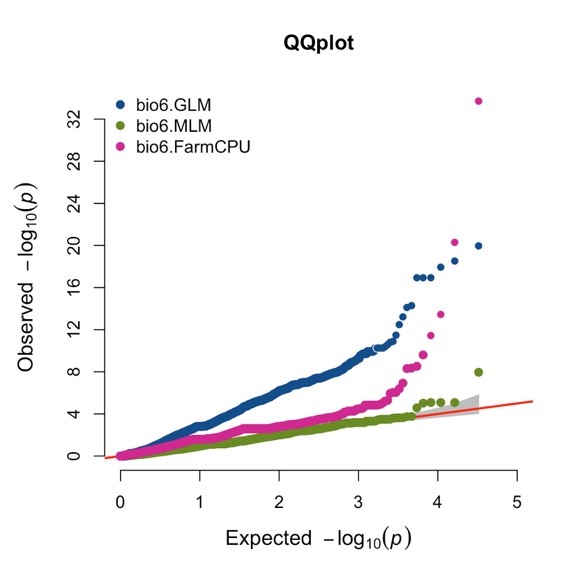

SI Figure 1. Quantile-Quantile plots for each environmental variable included in the EAA.

(continued)

(SI Figure 1. continued)

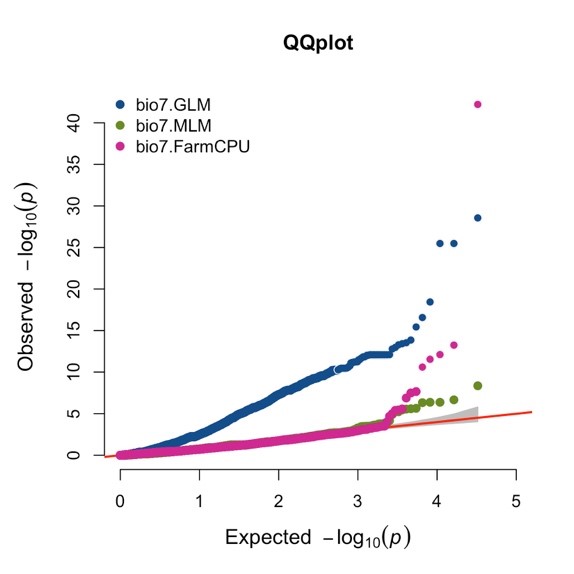

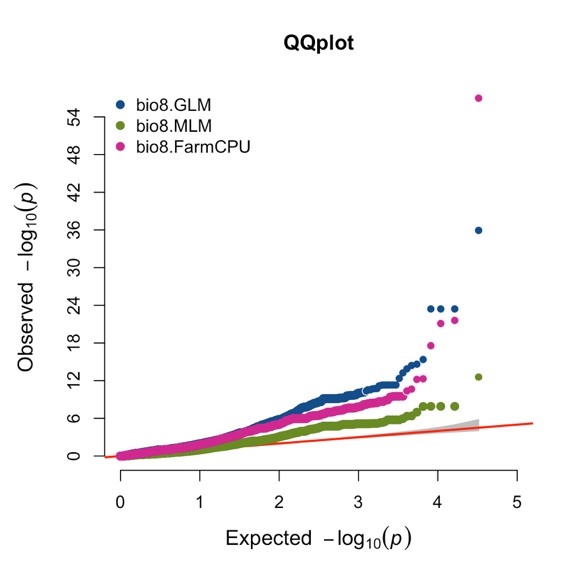

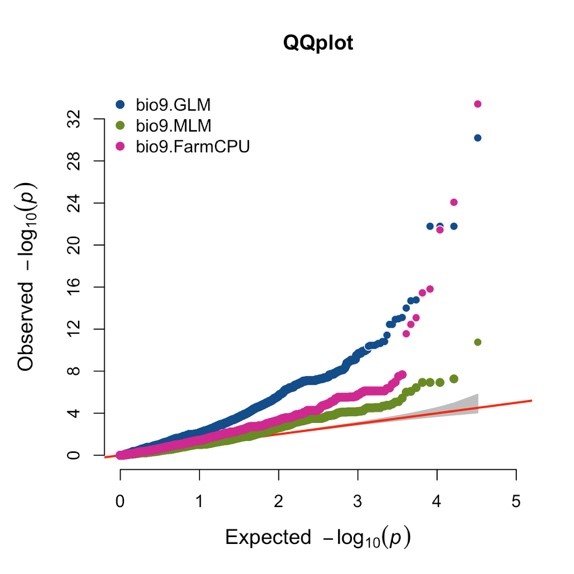

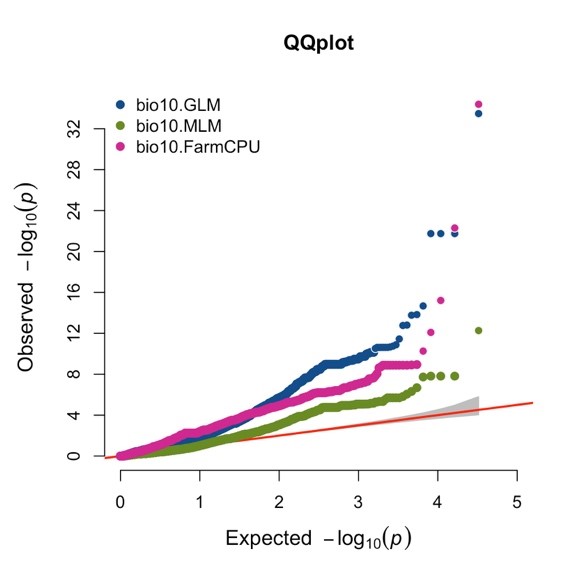

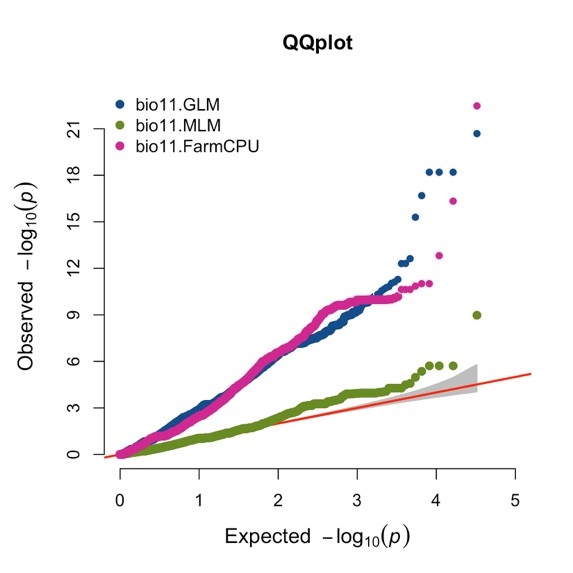

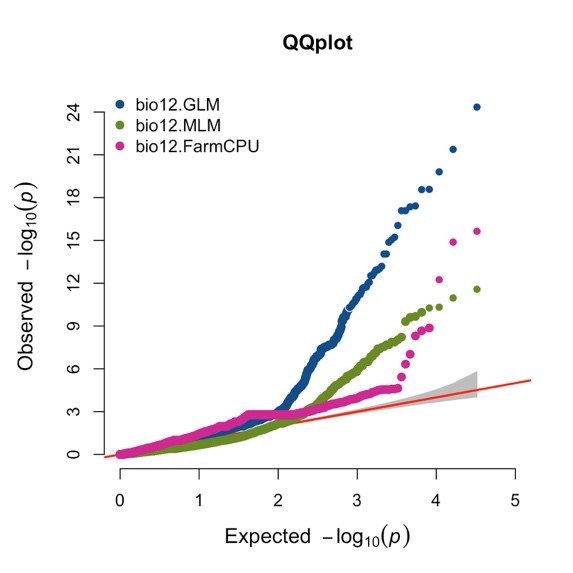

(continued)

(SI Figure 1. continued)

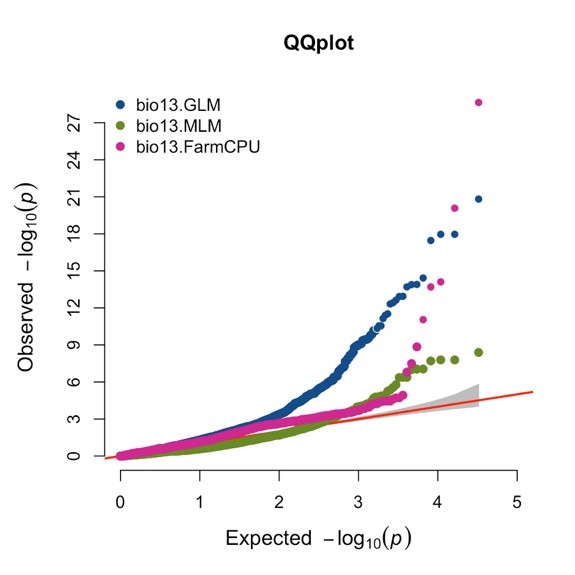

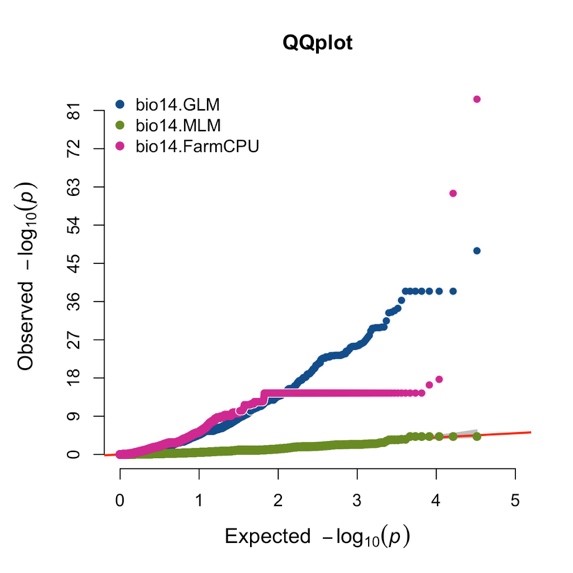

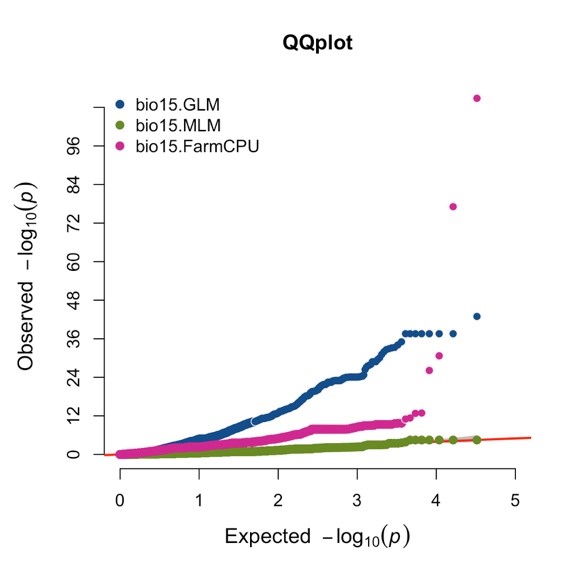

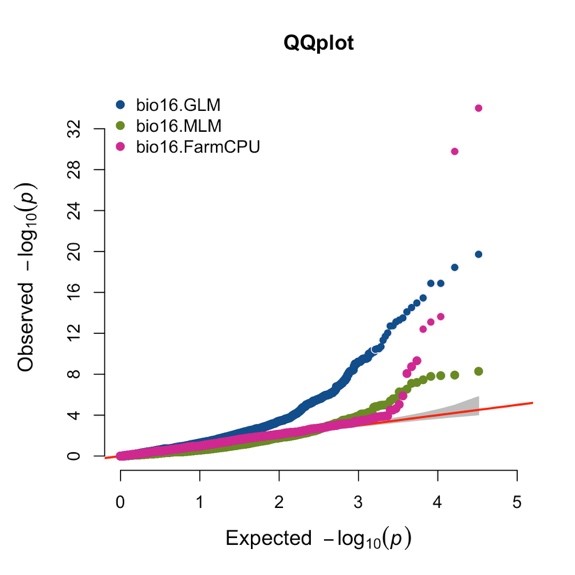

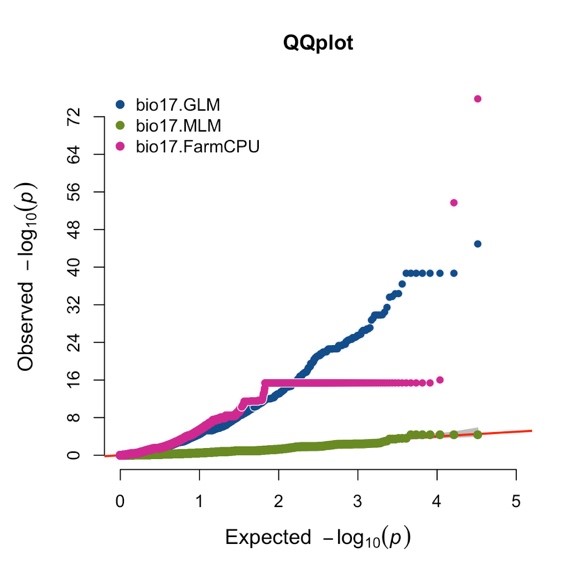

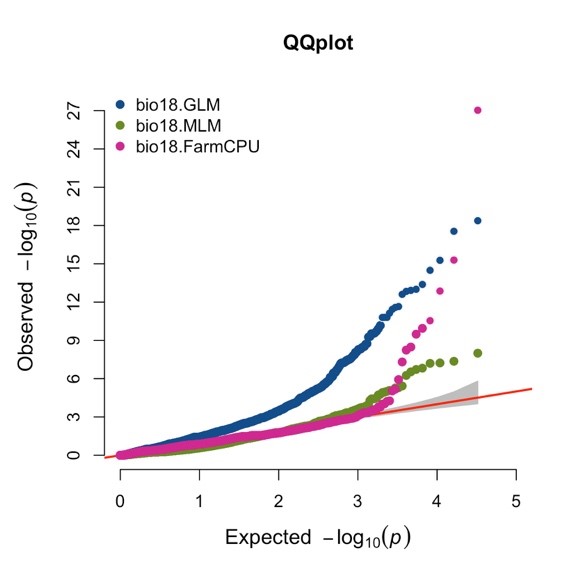

(continued)

(SI Figure 1. continued)

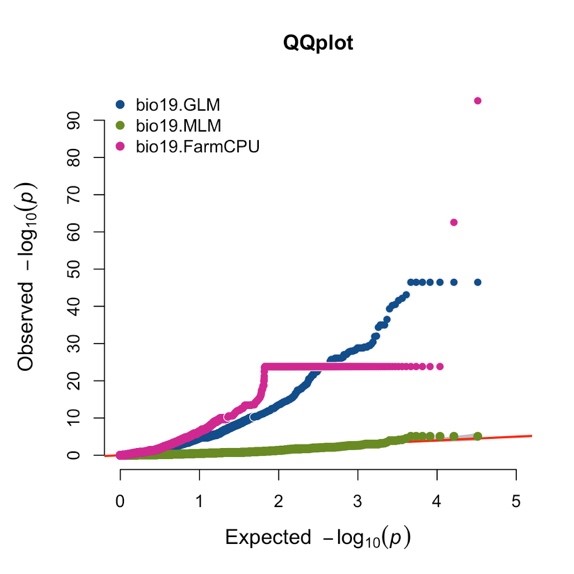

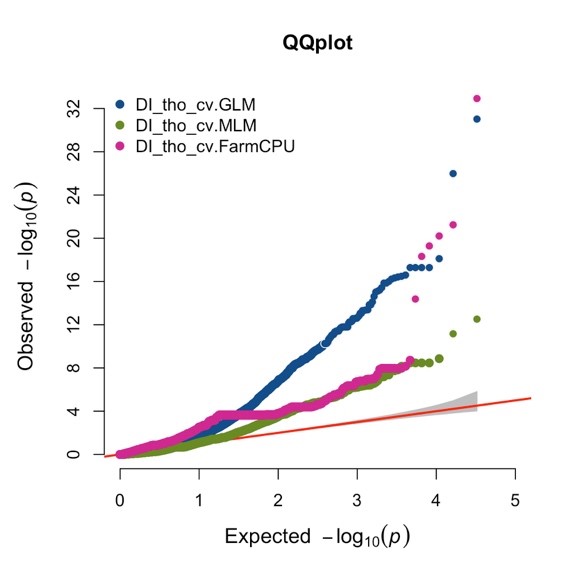

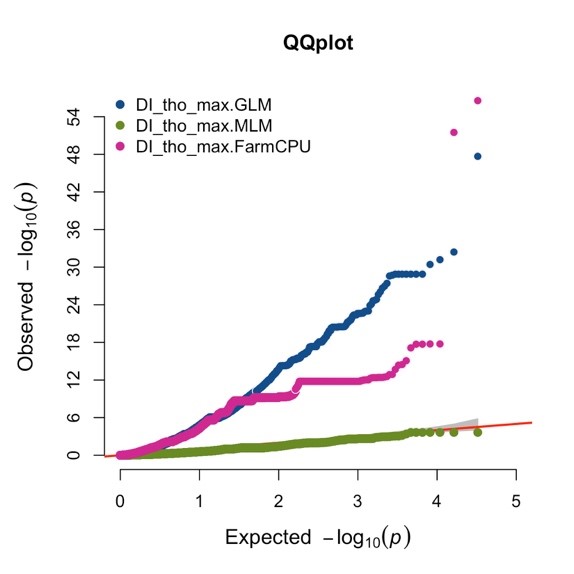

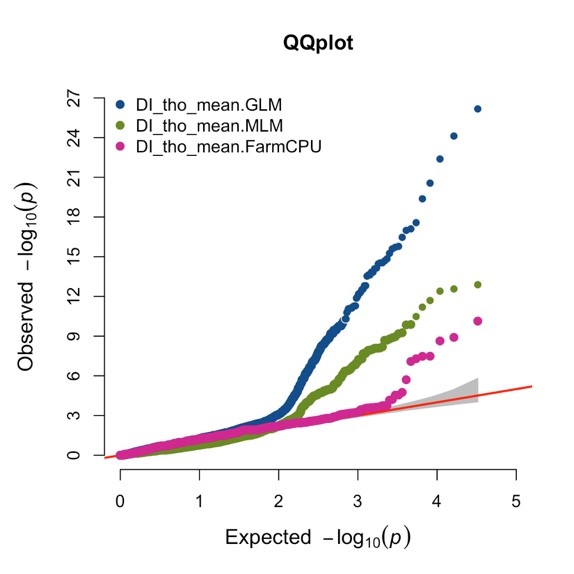

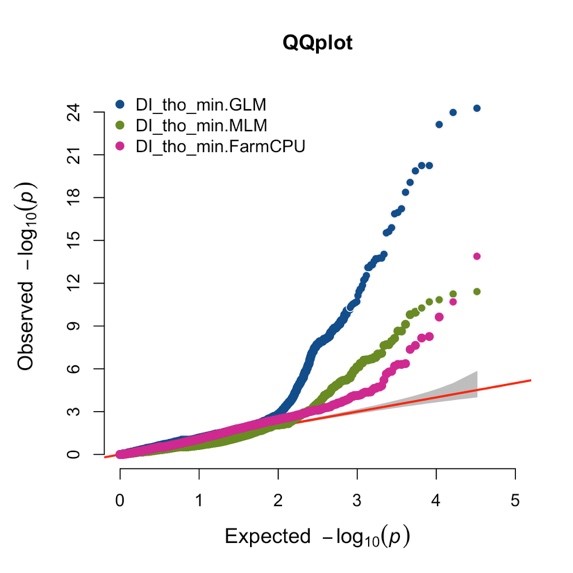

(continued)

(SI Figure 1. continued)

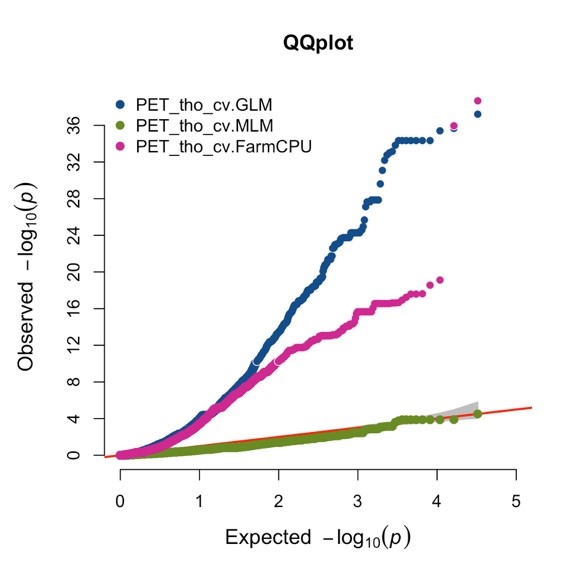

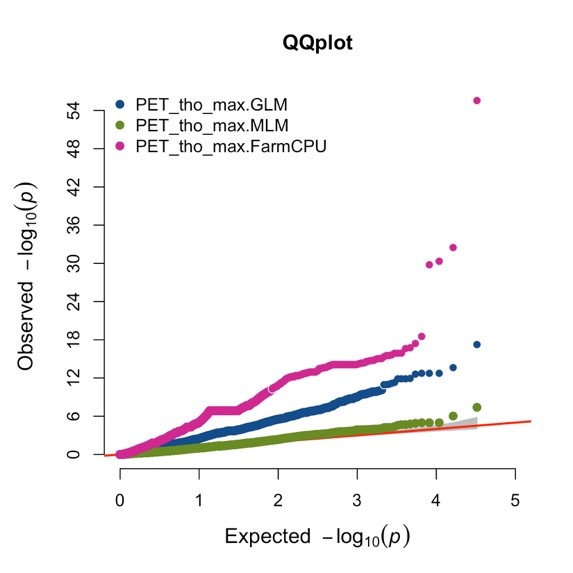

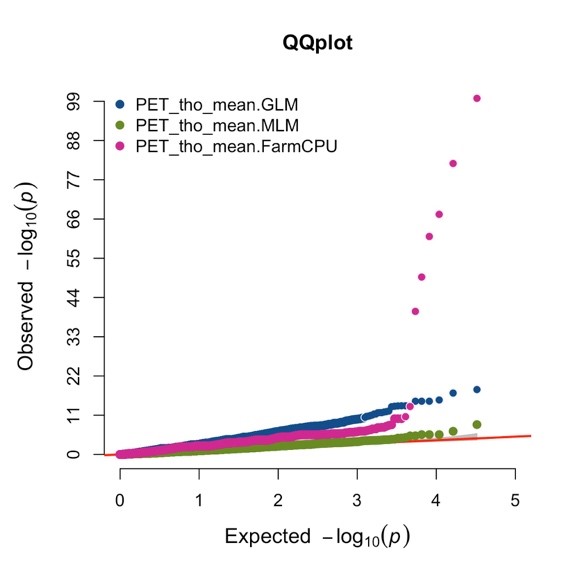

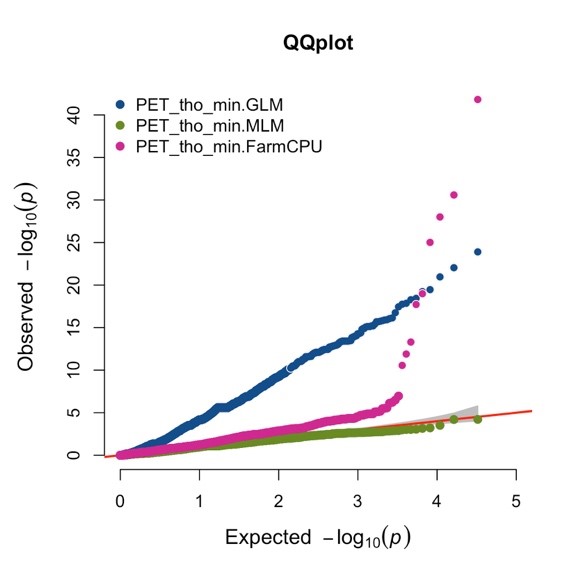

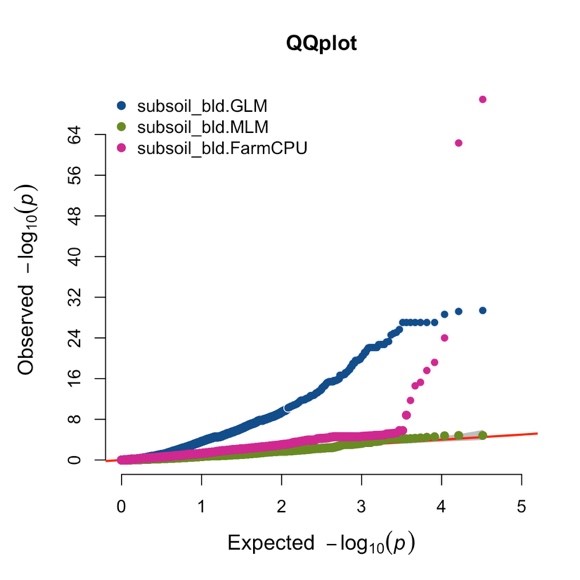

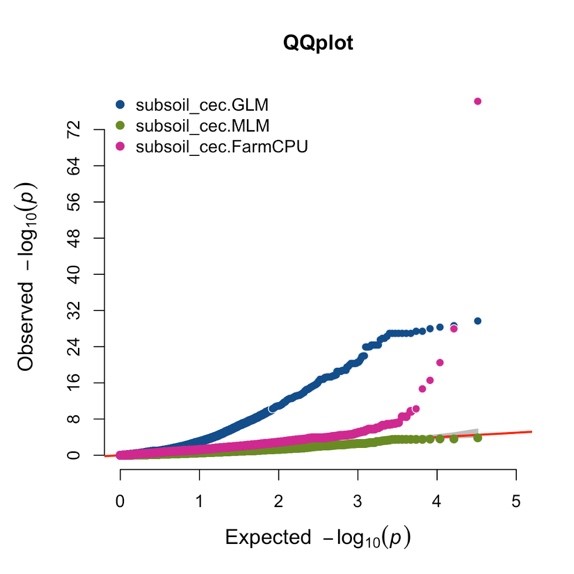

(continued)

(SI Figure 1. continued)

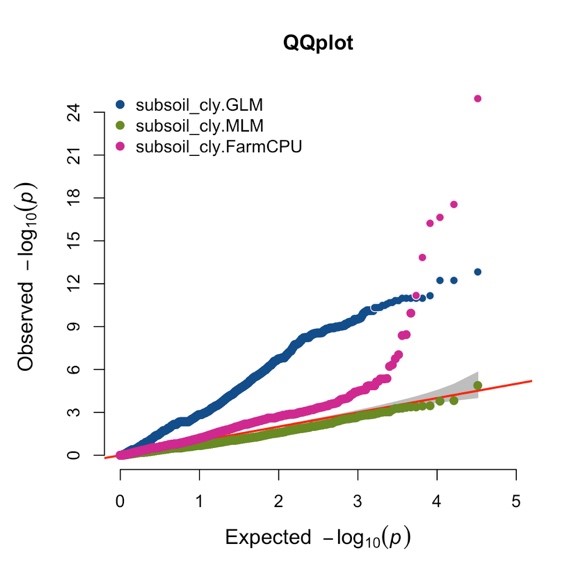

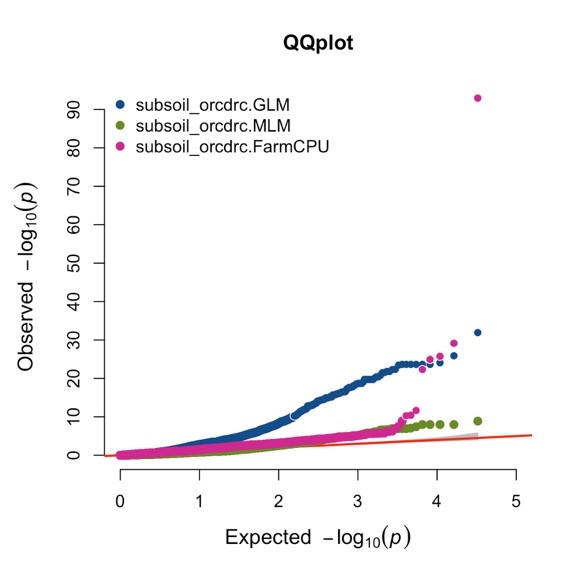

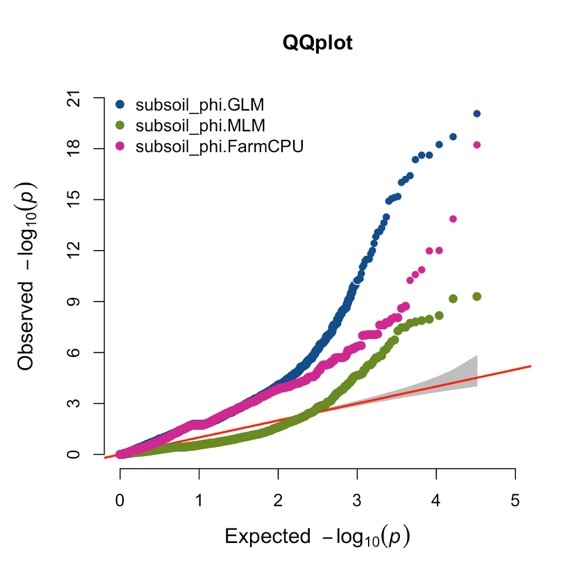

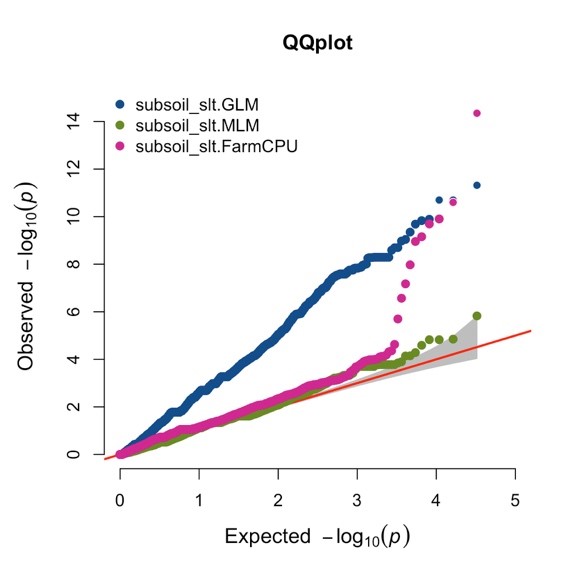

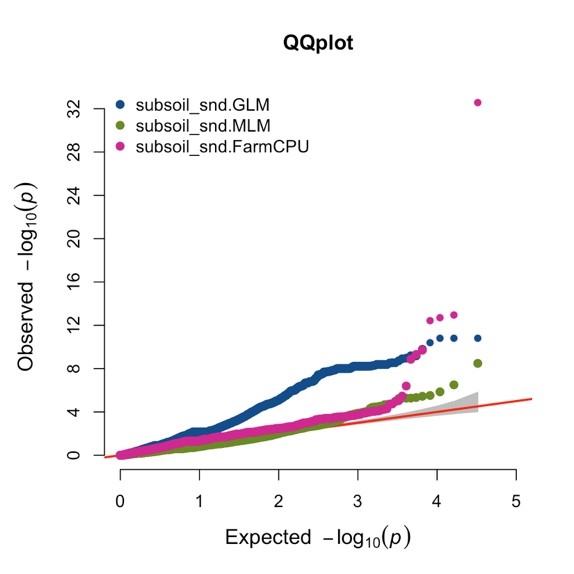

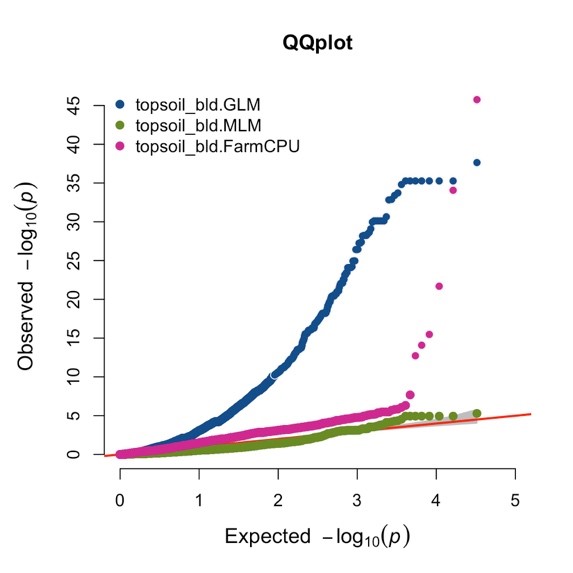

(continued)

(SI Figure 1. continued)

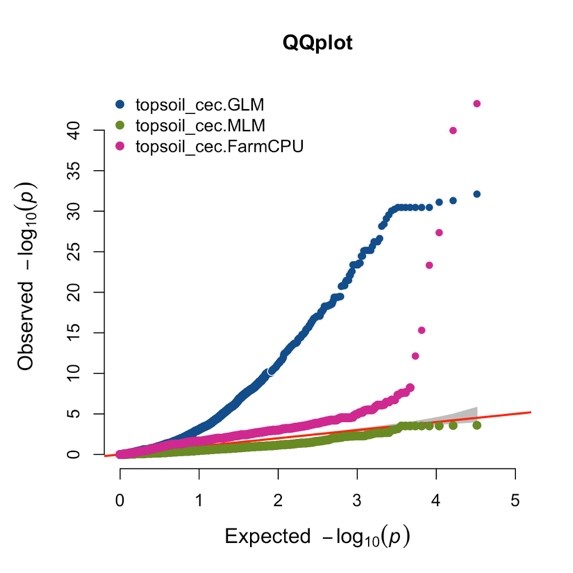

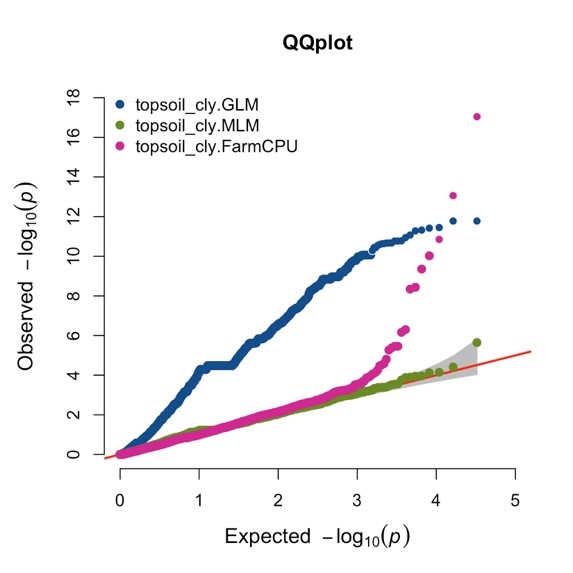

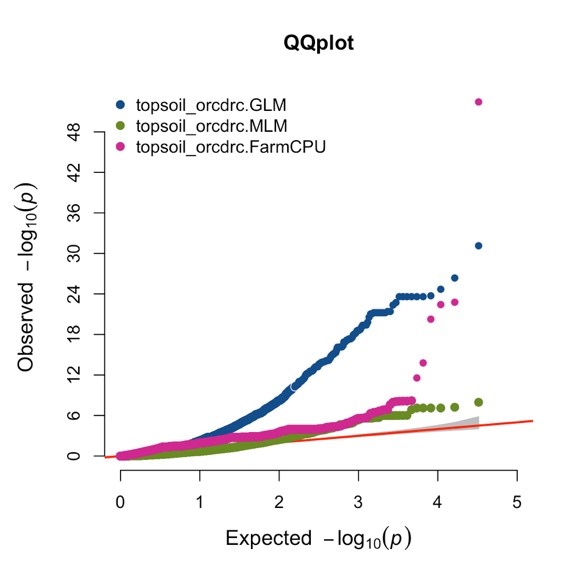

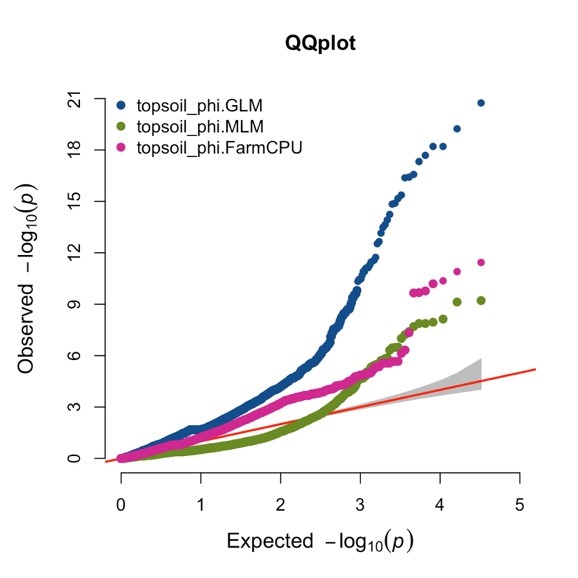

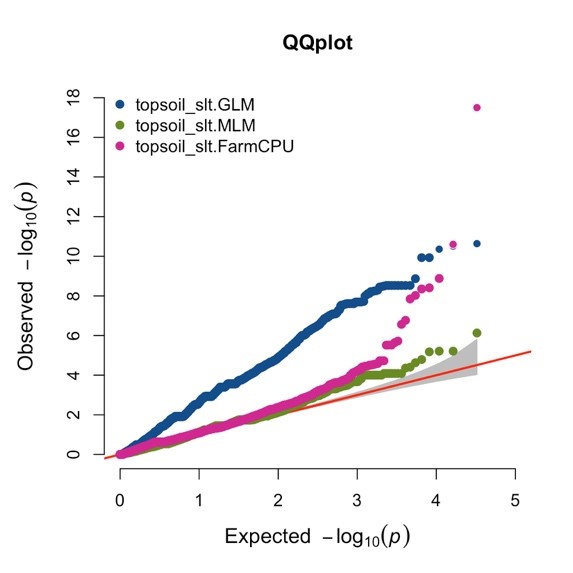

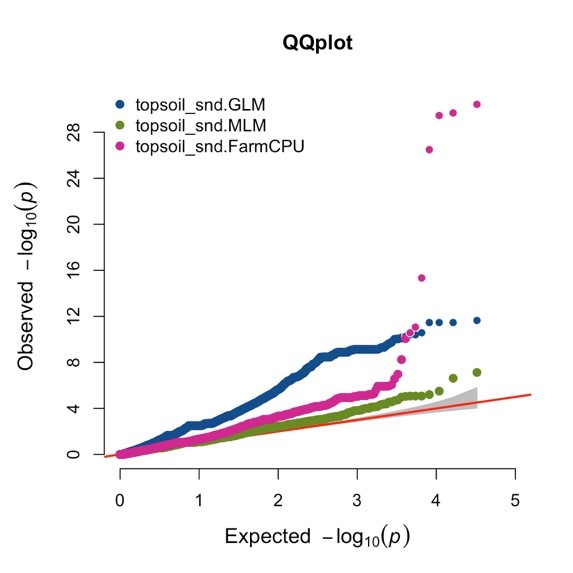
